# Genotype by Environment Interaction influenced the content of crude protein and total amino acids in lentil varieties

**DOI:** 10.1101/2025.08.25.672094

**Authors:** Alex Kumi Frimpong, Anastasiya Kuhalskaya, Lorenzo Rocchetti, Giacomo Conti, Asia Cicconi, Chiara Santamarina, Francesca Francioni, Benedetta Fanesi, Tania Gioia, Stefania Marzario, Alice Pieri, Giulia Frascarelli, Elisa Bellucci, Laura Nanni, Deborah Pacetti, Roberto Papa, Elena Bitocchi, Valerio Di Vittori

## Abstract

Lentil (*Lens culinaris* Medik) is a globally important grain legume valued for its high protein content and nutritional quality. However, the genetic and environmental factors influencing protein and amino acid composition, particularly the role of genotype × environment interaction (GEI), remain partially uncovered. This study evaluated 15 diverse lentil genotypes across seven agronomic trials spanning multiple years, sowing seasons, and locations to assess the effects of genotype, environment, and GEI on crude protein (CP), crude protein yield (CPY), total amino acids (TAA), and total amino acid yield (TAAY). Advanced statistical models such as AMMI, GGE, and WAASB were employed to dissect the contributions of genetic and environmental components and to identify stable, high-performing genotypes. Results revealed significant phenotypic variation for CP, Amino Acids (AA), and TAA among genotypes. Seasonal variation, especially between autumn and spring sowings, was the primary environmental driver of GEI for both CP and CPY. Notably, some landraces (PI_431710_LSP, PI_431739_LSP, PI_431753_LSP, IG_1959) demonstrated both high productivity and stability across environments, while others excelled in specific mega-environments identified through GGE analysis. Our findings emphasize the importance of integrating into lentil breeding programs, stability and adaptability, and a more comprehensive approach to measure the yield (e.g., CPY, MegaJoule, ammino acid composition and TAAY), which takes into account the quality and the effective energetic production of a crop. From this perspective, we highlight landraces as valuable sources of genetic diversity for improving yield. This work provides a foundation for targeted breeding strategies aimed at developing lentil varieties with enhanced protein content, balanced amino acid profiles, and resilience to environmental variability

## INTRODUCTION

Lentil (*Lens culinaris* Medik) is an important grain legume cultivated globally due to its high nutritional value and adaptability to various agro-climatic conditions (Khazaei et al., 2019; Kumar et al., 2016). It plays an important role in global food security, especially in regions where a plant-based diet is a staple (Stagnari et al., 2017). It is a diploid (2n = 14), self-pollinating annual crop, and originated in the Middle East (Cokkizgin & Shtaya, 2013). Lentil is known to be one of the earliest domesticated plants cultivated, tracing back to about 10,000 years (Guerra-Garcia et al., 2022; Liber et al., 2021). Currently, lentils are cultivated in more than 52 countries, covering approximately 5.58 million hectares, with an annual production of 6.65 million metric tons (FAOSTAT, 2022). Canada and India are the leading producers, together contributing over 60% of the total global production (Khazaei et al., 2019). The crop’s ability to fix atmospheric nitrogen through symbiosis with *Rhizobium* spp. can contribute to enhancing soil fertility, decreasing the reliance on synthetic fertilizers, and promoting sustainable agricultural practices (Siddique et al., 2023; Stagnari et al., 2017). Furthermore, lentils exhibit moderate drought tolerance, making them suitable for cultivation in arid and semi-arid regions (Kumar et al., 2016; Subedi et al., 2021). Cultivated lentils are highly valued for their rich protein content, with an average of 26% of dry biomass (Dhull et al., 2023; Iqbal et al., 2006; Joshi et al., 2017). Earlier studies performed in various regions of the world have recorded wide variabilities in the crude protein content, ranging between 20% to 36% of dry biomass (Karaköy et al., 2012; Tahir et al., 2011; Wang & Daun, 2006). In addition to protein, lentils provide essential amino acids, fiber, micronutrients (iron, zinc, and folate), and bioactive compounds with health-promoting properties (Carbonaro & Nucara, 2022; Tessari et al., 2016; Wu, 2013). However, like most legumes, lentils are low in sulfur-containing amino acids, such as methionine and cysteine, which can be resolved by dietary complementation with cereals (Margier et al., 2018; van Vliet et al., 2015). Several QTLs are thought to control the genetic inheritance of the nutritional trait, and due to its complex nature (i.e., quantitative trait), the environmental conditions can significantly contribute to the total phenotypic variation of the trait (Collard et al., 2005). Genes and QTLs associated to total protein contents and related traits have been identified (Upadhyaya et al., 2016). Despite their nutritional and agricultural benefits, lentil production encounters challenges related to genotype-by-environment (G × E) interactions, which significantly impact yield, protein content, and amino acid composition (Acuña & Wade, 2013; Annicchiarico, 2002; Aruna et al., 2016; Subedi et al., 2021). Environmental factors such as temperature, precipitation, and soil fertility influence seed composition, often resulting in significant variation across environments (Bansal et al., 2023; Chen et al., 2022; M. S. Erskine, 1985; Zeroual et al., 2023). The identification of genotypes with superior adaptability and nutritional traits relies on robust agronomic characterization of genetic resources. The INCREASE project (Bellucci et al., 2021) exemplifies this effort by integrating genomic, phenotypic, and phenomic data across major legumes such as common bean and lentil (Cortinovis et al., 2021; Guerra-Garcia et al., 2022). Moreover, understanding the genetic basis of protein content and other nutritional profiles is vital for developing improved cultivars with enhanced nutritional traits (Johnson et al., 2024; Margier et al., 2018; Salaria et al., 2022).

Statistical models have been established to assess G × E interaction and genotype performance based on multi-environment trials. The Additive Main Effects and Multiplicative Interaction (AMMI) model partitions genetic and environmental effects and provides insights into both the stability and adaptability of genotypes (Gauch Jr., 2006; Gauch Jr. et al., 2008; Yan et al., 2007). Similarly, the Genotype plus Genotype-by-Environment Interaction (GGE) biplot method evaluates both genetic and environmental effects on trait performance (Crossa et al., 1999; Yan et al., 2000). More recently, the Weighted Average of Absolute Scores (WAASB) index has been employed to quantify genotype stability across diverse environments (Olivoto et al., 2019; Piepho, 1994). Integrating these statistical approaches enables plant breeders to select genotypes with superior nutritional attributes while ensuring adaptability to different environmental conditions. In lentils and other pulses, protein content and yield tend to be negatively correlated; that is, a higher yield can be associated with a lower protein content. Thus, a simultaneous positive selection for high yield and protein content can be challenging for breeders (W. Erskine et al., 2003; Tziouvalekas et al., 2022; Vlachostergios et al., 2021). Given the increasing demand for plant-based protein sources and the significance of worldwide nutritional security, it is crucial to identify lentil genotypes displaying high and stable protein content across multiple environments (Hang et al., 2022; Pugliese et al., 2024). Therefore, this study aims to evaluate the effects of genotype, environment, and G × E interactions on crude protein content, crude protein yield, total amino acid composition, and amino acid yield in lentils. By employing AMMI, GGE, and WAASB models, we identified high-performing and stable genotypes that can serve as promising candidates for future breeding programs.

## MATERIALS AND METHODS

### Plant Material

A total of 324 Single Seed Descent (SSD) lentil genotypes from the AGILE project (*Application of Genomic Innovation in the Lentil Economy, KnowPulse*, led by the University of Saskatoon, Canada), were previously tested in a field trial carried out in 2016-2017 season in Metaponto (Italy), in collaboration with the University of Basilicata (Wright et al., 2021). The lentil panel mostly includes landraces from 45 different countries. Out of the entire panel, we initially selected 30 lentil lines that showed the highest yield in the Metaponto field trial (Rocchetti et al., 2025; Wright et al., 2021). We added 11 Italian landraces previously collected from local farmers and seed companies, three Italian cultivars provided by AgroService ISEA, and two French cultivars provided by AgriObtention, reaching a total number of 46 lentil lines **Supplementary Table S1** (Rocchetti et al., 2025).

The 46 lines were evaluated across seven agronomic trials that have been carried out in three years (2019, 2020, 2021), two sowing seasons (autumn and spring) and two Italian localities, that are the Research Centre for Cereal and Industrial Crops (CREA-CI) experimental station of Osimo (Ancona, latitude, 43° 45’ 04”, longitude, 13° 49’ 98”) and the experimental station of Metaponto (University of Potenza, Matera, latitude 40° 23’ 24’’, longitude 16° 46’ 48’’) (**Table 1**).

**Table 1.** List of seven field trials (environments) given by the combination of localities, sowing season, and years of cultivation.

| Locality | Metaponto |  | Osimo |  |  |  |  |
| --- | --- | --- | --- | --- | --- | --- | --- |
| Sowing | Autumn |  | Autumn |  |  | Spring |  |
| Year | 2019 | 2020 | 2019 | 2020 | 2021 | 2020 | 2021 |
| Name of the trial | Metaponto<br>autumn 2019 | Metaponto<br>autumn 2020 | Osimo<br>autumn<br>2019 | Osimo<br>autumn<br>2020 | Osimo<br>autumn<br>2021 | Osimo<br>spring<br>2020 | Osimo<br>spring<br>2019 |

Those trials allowed further testing of the agronomical value of the selected material.

The following environmental variables were recorded for the Metaponto and Osimo locations for the three years (2019, 2021, and 2022) and sowing seasons (Autumn and Spring): average temperature (°C), max temperature (°C), minimum temperature (°C), and rainfall (mm) (**Supplementary Figure S1**). Field trials were conducted using a Randomized Complete Block Design (RCBD) with three replicates of 10m^2^ plot size, and six rows with 0.15 m distance between rows per plot. Sowing was conducted during December for the autumn sowing and during March for the spring sowing. For details, see (Rocchetti et al., 2025).

Out of the 46 lines, we selected the most promising 15 lines, for which seeds were also obtained in at least two replicates across all seven environments to evaluate the nutritional quality (**Table 2**).

**Table 2.** List of genotypes for which at least two replicates (seed material for nutritional analysis) were available at the seven environments evaluated in this study.

| **Genotypes** | **Donor** | **Origin** | **Biological Status** |
| --- | --- | --- | --- |
| PI_612875_LSP | AGILE/UNIBAS | Syria | Breeding material |
| W6_27760_LSP | AGILE/UNIBAS | USDA | Breeding material |
| PI_533693_LSP | AGILE/UNIBAS | Spain | Cultivar |
| Anicia | Agri Obtentions | France | Cultivar |
| Flora | Agri Obtentions | France | Cultivar |
| PI_298122_LSP | AGILE/UNIBAS | France | Cultivated material |
| PI_299351_LSP | AGILE/UNIBAS | Chile | Cultivated material |
| IG_1959 | AGILE/UNIBAS | Ethiopia | Landrace |
| PI_426778_LSP | AGILE/UNIBAS | Pakistan | Landrace |
| PI_431663_LSP | AGILE/UNIBAS | Iran | Landrace |
| PI_431710_LSP | AGILE/UNIBAS | Iran | Landrace |
| PI_431728_LSP | AGILE/UNIBAS | Iran | Landrace |
| PI_431739_LSP | AGILE/UNIBAS | Iran | Landrace |
| PI_431753_LSP | AGILE/UNIBAS | Iran | Landrace |
| PI_432033_LSP | AGILE/UNIBAS | Iran | Landrace |

### Crude Protein Determination

A total of 284 samples, which are the single replicates of the genotypes across different environments, were analyzed to determine the crude protein content. Dry seeds from each sample were ground into a fine powder using liquid nitrogen with a prechilled mortar and pestle to ensure minimal degradation. Approximately 70 mg of the powdered sample was aliquoted into pre-labeled tin foil cups (Leco 502-186) and sealed into teardrop-shaped pockets for analysis.

Nitrogen (N), carbon (C), and hydrogen (H) contents were determined using the Dumas combustion method with an elemental analyzer (TruSpec Micro, LECO Corp., St. Joseph, MI, USA). Nitrogen content was used to quantify the total Crude protein (CP) for each analyzed sample, based on the Jones factor (N × 6.25), which is a widely accepted conversion factor for protein determination in dry biomass (Mossé & Baudet, 1983; Salo-väänänen & Koivistoinen, 1996).

The crude protein yield (CPY) was estimated as follows:

CPY (kg ha□¹) = Crude Protein Content (%) × Seed Yield (kg ha□¹)

### Amino Acid (AA) Extraction and Determination

For amino acid quantification, 20 mg of freeze-dried sample was hydrolyzed with 6 M HCl (500 µL) at 110 °C for 24 hours. The hydrolysate was then dried under a speed vacuum for 1 hour, reconstituted with 0.1 M HCl (500 µL), and centrifuged (6000 rpm, 5 minutes, 4°C). A ten-fold dilution of the supernatant was prepared, and 50 µL of the diluted extract was combined with DL-norvaline (50 µL of 0.05 mg/mL in 0.1 M HCl) as an internal standard. The mixture was dried again and derivatized using the AccQ·Tag™ Ultra Derivatization Kit (Waters Corporation, Milford, MA, USA). Subsequently, 1 µL of each derivatized sample was injected into a UPLC (Waters, USA) system equipped with a photodiode array (PDA) detector and an AccQ Tag Ultra Column (2.1 x 100 mm, 1.7 µm). Ultra High-Performance Liquid Chromatography (UHPLC) data were normalized to account for run-to-run variability across days, and the resulting normalized matrix was used for amino acid quantification. Standard calibration curves were prepared using a mixture of amino acids (1 pM to 25 pM in 0.1 M HCl) to represent each concentration level. Essential amino acids (EAA) identified included Histidine (His), Threonine (Thr), Lysine (Lys), Tyrosine (Tyr), Methionine (Met), Valine (Val), Leucine (Leu), Isoleucine (Ile), and Phenylalanine (Phe). Non-essential amino acids (NEAA) comprised Alanine (Ala), Arginine (Arg), Aspartic acid (Asp), Cysteine (Cys), Glutamic acid (Glu), Glycine (Gly), Proline (Pro), and Serine (Ser). The total amino acid yield (TAAY) was calculated as follows:

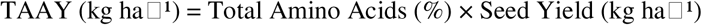

Data for crude protein content, total amino acids (TAA), essential amino acids (EAA), crude protein yield, TAA yield, and the 17 detected amino acids are presented in **Supplementary Table S2**.

### Analysis of variance and trait heritability

The phenotypic variability and heritability for nutritional traits in the 15 genotypes grown under different environments have been investigated. A one-way ANOVA followed by post-hoc Tukey’s Honestly Significant Differences (Tukey’s HSD) analysis was performed to identify significant differences among the sample means. For data visualization, boxplots have been generated by using the online software Metaboanalyst 5.0 (Pang et al., 2021). Considering the linear model (1), the phenotypic value of a trait of any individual in a given environment can be obtained based on overall mean (μ), genetic (G), environmental (E), genotype by environment interaction (GEI) effects, and *e*, which is the residual effect within each environment, assuming it is normally distributed;

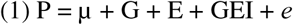

The Residual Maximum Likelihood (RELM) was used to estimate variance components, which are G, E, and GEI effects, and the associated standard errors, as random factors. Broad-sense heritability (H^2^) was estimated following the variance partition, using the following formula (2):

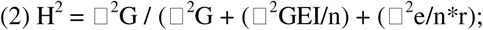

where ^2^G represents the genetic variance, ^2^GEI the variance of genotype-environment interaction, *n* the number of environments, *r* is the number of replicates in each environment, and e is the residual error. Based on model (1), we performed ANOVA to test the significance of G, E, and GEI effects, as fixed factors. The environment was dissected into locality (L) and sowing season (S) effects. Statistically significant differences among genotypes, localities, and seasons were determined by using Tukey’s test, at the probability level of p < 0.05. Two multivariate analyses were conducted: i) a principal component analysis (PCA) was carried out based on the mean value of each trait in each environment to investigate the relationships among the different genotypes by JMP^®^ Pro, Version 18.0.2 (*JMP®*, 1989); ii) a heatmap, representing abundances of each trait by false color imaging on the log_10_-transformed data across the genotypes, combined with two-dimensional hierarchical clustering, performed with the hclust function in the package stat implemented in Metaboanalyst ver.5.0 (Pang et al., 2021). Similarity was computed through Euclidean distance, employing the Ward D clustering algorithm, which aims to cluster by minimizing the sum of squares of any two clusters. To identify key traits driving variation among the amino acids, we used PLS-DA to generate a Variable Importance in Projection (VIP) plot, highlighting the most influential nutritional compounds based on their relative concentrations (**Supplementary figure S2**). One-way analysis of variance (ANOVA) was computed to compare genotypes grouped by their biological status (landraces *versus* cultivar and breeding materials) for the nutritional traits considered. Significant differences among groups were tested by applying Tukey’s test (p < 0.05).

### Correlation analysis

Pearson correlation analysis was performed among the mean values of the amino acids, the total amino acid value, and the crude protein content, using JMP Pro version 18.0.2 (*JMP®*, 1989).

### Additive main effect and multiplicative interaction (AMMI)

Nitrogen content data from all the test environments were subjected to the Additive Main Effect and Multiplicative Interaction (AMMI) (Olivoto et al., 2019). AMMI combines ANOVA and SVD to break down genotype × environment interactions and uses biplots to visually interpret these effects. AMMI analysis was employed to evaluate the GEI effect, identify specific positive genotype-environment interactions, or determine stable genotypes. This is achieved by dissecting GEI into principal components (i.e., PC in a Principal Component Analysis. The first Principal Component Axis (PCA1) and the first two Principal Component Axes (PCA1 and PCA2) were used to generate AMMI biplots (PCA1 vs. PCA2).

### Genotype plus genotype by environment interaction (GGE)

The Genotype plus Genotype-environment interaction (GGE) model (Yan et al., 2000) considers the joint effects of the genotypic main effect, together with GEI. This model was used to investigate trait productivity and to identify similarity among environmental performances, based on the genotype rankings, and the best genotypes performing in each environment. Results are presented using GGE biplots, which approximate overall performance (G + GEI).

### Stability with WAASB (Weighted Average of Absolute Scores)

The stability index WAASB (Weighted Average of Absolute Scores) developed by Olivotto et al. (2019) was used to model the genotype performance versus stability across environments. This model applies the SVD (Singular Value Decomposition) of the BLUP matrix for the GEI effect generated by Linear Mixed Model (LMM). The genotype with the lowest WAASB value is considered the most stable: that is, the one that deviates the least from the average performance across environments. Results were reported using scatter plots for each trait, representing the phenotypic values of each genotype (abscissa) and the WAASB index (ordinate).

## RESULTS

### Phenotypic variation for nutritional traits

The total amino acid (TAA) content has been investigated in the seeds from the selected lentil accessions previously grown across different environments and years of trial. The amount of 17 amino acids was quantified by high-throughput UHPLC (**Figure 1**). CP, CPY, TAA, and TAAY have been quantified for each genotype, considering all replicates in each environment (**Supplementary Figure S3**). A higher phenotypic variability can be observed among genotypes when considering CP, compared to TAA. The average values across replicates and environments for each genotype, and summary statistics, are provided for the nutritional traits that are under investigation (**Table 3**). CP content ranged from 24.31% to 35.90%, with a mean of 29.27%, higher than the typical values reported in the literature for lentil. Similarly, variation was observed in both TAA and total essential amino acid (TEAA) contents, which ranged from 8.57 to 19.80 mg/100 mg (mean: 14.26 mg/100 mg) and from 4.60 to 8.00 mg/100 mg (mean: 6.05 mg/100 mg), respectively (**Table 3**).

**Figure 1.**
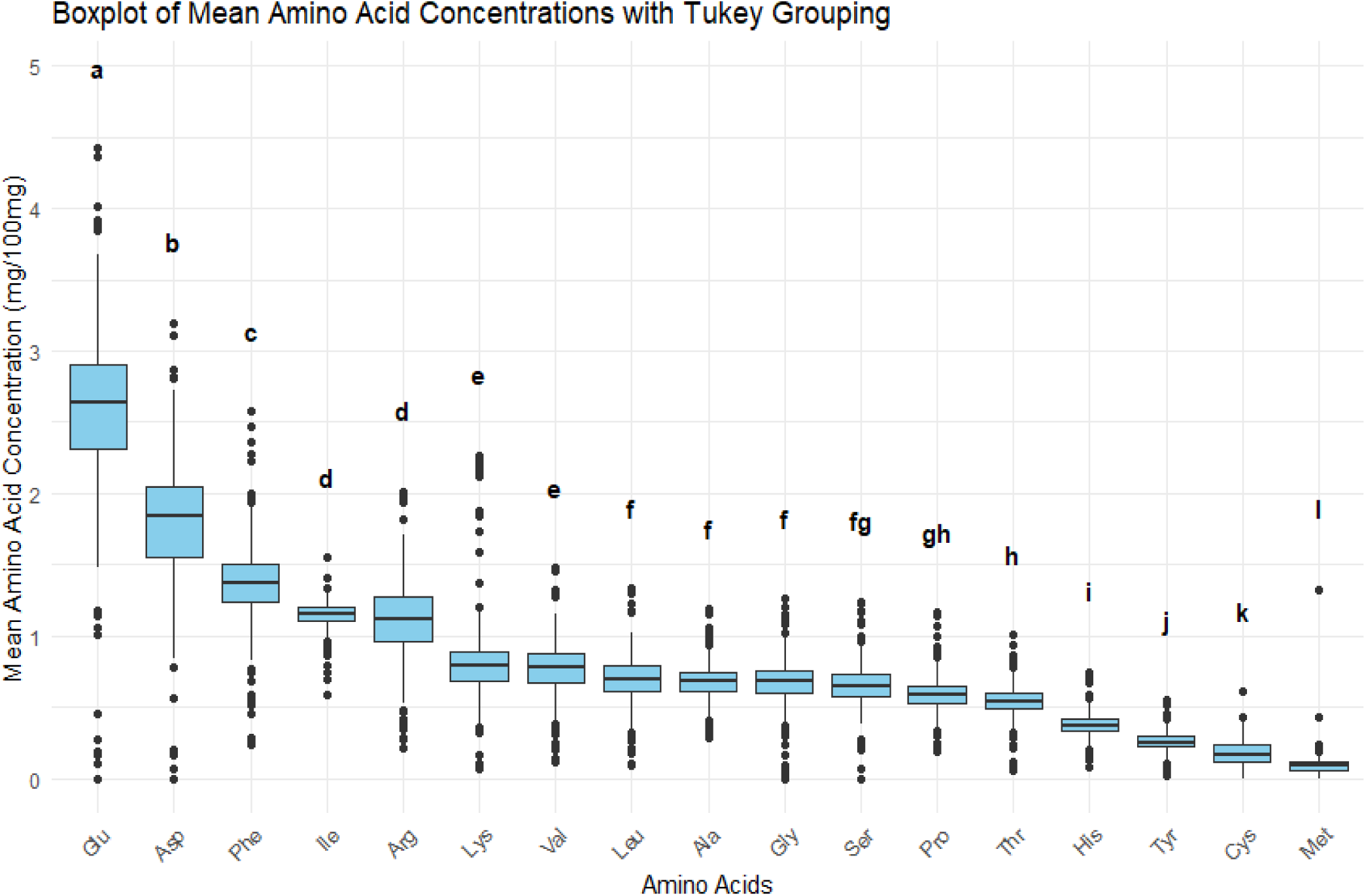
Representation of the individual amino acid profiles (nutritional quality) across 15 lentil genotypes. A Box plot of the mean and range of all amino acidcontent for 15 lentil genotypes averaged across localities and years.

**Table 3:** Average values across replicates and environments for crude protein, crude protein yield, and amino acid composition for 15 selected lentil genotypes. Minimum, maximum values, standard deviation, coefficient of variation, and value range are provided.

| **Trait** | **Mean** | **Min** | **Max** | **Std Dev** | **CV** | **Range** |
| --- | --- | --- | --- | --- | --- | --- |
| Crude protein (%) | 29.27 | 24.31 | 35.90 | 2.17 | 7.45 | 11.60 |
| Protein yield (kg/ha) | 156.95 | 6.65 | 514.43 | 114.14 | 73.07 | 507.79 |
| TAA (mg/100mg) | 14.26 | 8.57 | 19.85 | 1.95 | 13.77 | 11.27 |
| Amino Acid yield (kg/ha) | 76.66 | 2.90 | 259.42 | 57.62 | 75.53 | 256.51 |
| TEAA (mg/100mg) | 6.05 | 4.64 | 8.00 | 0.60 | 10.03 | 3.35 |
| **ESSENTIAL AMINO ACIDS (mg/100mg)** |  |  |  |  |  |  |
| His | 0.38 | 0.26 | 0.55 | 0.06 | 15.31 | 0.29 |
| Thr | 0.54 | 0.37 | 0.73 | 0.07 | 13.29 | 0.36 |
| Val | 0.77 | 0.51 | 1.15 | 0.12 | 15.65 | 0.64 |
| Lys | 0.84 | 0.51 | 1.60 | 0.21 | 25.52 | 1.09 |
| Ile | 1.14 | 0.91 | 1.31 | 0.07 | 5.97 | 0.40 |
| Leu | 0.69 | 0.44 | 1.06 | 0.11 | 15.99 | 0.61 |
| Phe | 1.36 | 0.95 | 1.94 | 0.19 | 14.23 | 0.98 |
| Met | 0.09 | 0.00 | 0.73 | 0.08 | 85.82 | 0.73 |
| **NON-ESSENTIAL AMINO ACIDS (mg/100mg)** |  |  |  |  |  |  |
| Ser | 0.65 | 0.00 | 0.96 | 0.13 | 19.80 | 0.96 |
| Arg | 1.12 | 0.70 | 1.62 | 0.19 | 17.00 | 0.91 |
| Gly | 0.68 | 0.00 | 1.14 | 0.13 | 19.44 | 1.14 |
| Asp | 1.78 | 0.00 | 2.83 | 0.39 | 21.81 | 2.83 |
| Glu | 2.57 | 0.00 | 3.90 | 0.48 | 18.89 | 3.90 |
| Ala | 0.68 | 0.50 | 0.92 | 0.08 | 12.48 | 0.43 |
| Tyr | 0.26 | 0.12 | 0.38 | 0.05 | 19.30 | 0.26 |
| Pro | 0.59 | 0.42 | 0.86 | 0.08 | 14.35 | 0.45 |
| Cys | 0.18 | 0.00 | 0.43 | 0.07 | 40.61 | 0.43 |
| **TOTAL SULFUR AMINO ACID (mg/100mg)** |  |  |  |  |  |  |
| TSAA(Met+Cys) | 0.23 | 0.00 | 0.63 | 0.09 | 38.94 | 0.63 |

With regards to the single amino acid (AA) content, the sulphur-containing amino acids (Met and Cys) are present in relatively low quantities, that is, 0.09mg/100mg and 0.18mg/100mg, respectively, as could be expected in lentils and legumes in general. However, variation can be observed among the traits, with TSAA content varying from 0mg/100mg (not detected) to 0.63mg/100mg (**Table 3**). High contents of Glu and Asp and intermediate levels of Phe, Ile, and Arg can be observed in the analyzed samples. (**Figure 1**, **Table 3**). Results for the two-dimensional hierarchical cluster analysis (2D-HCA) and relative heat map for the considered traits (mean values of all the replicates of each genotype) are provided in **Supplementary Figure S4**, supporting the presence of significant phenotypic variation among the analyzed samples.

### ANOVA, Heritability, and Correlation among the nutritional traits

Broad-sense heritability (H^2^) was estimated for 22 traits (**Table 4**). Focusing on each AA, H^2^ ranged from 0 (Lys, Met, and Ile) to 0.78 (Cys). The Asp and Glu AAs showed relatively lower heritability (H^2^ of 0.21 and 0.12, respectively) when compared to other AAs. Eight AAs (His, Arg, Ala, Pro, Cys, and Tyr) showed high heritability (H^2^ > 0.59), while four AAs (Ser, Gly, Val, and Leu) showed an intermediate estimated H^2^ (0.37 < H^2^ < 0.44). Also, except for TEAA, for which we estimated a lower heritability (H^2^ = 0.13), CP, TAA, CPY, and TAAY showed high heritability estimates (H^2^ = 0.78, 0.43, 0.66, and 0.59, respectively).

**Table 4:**
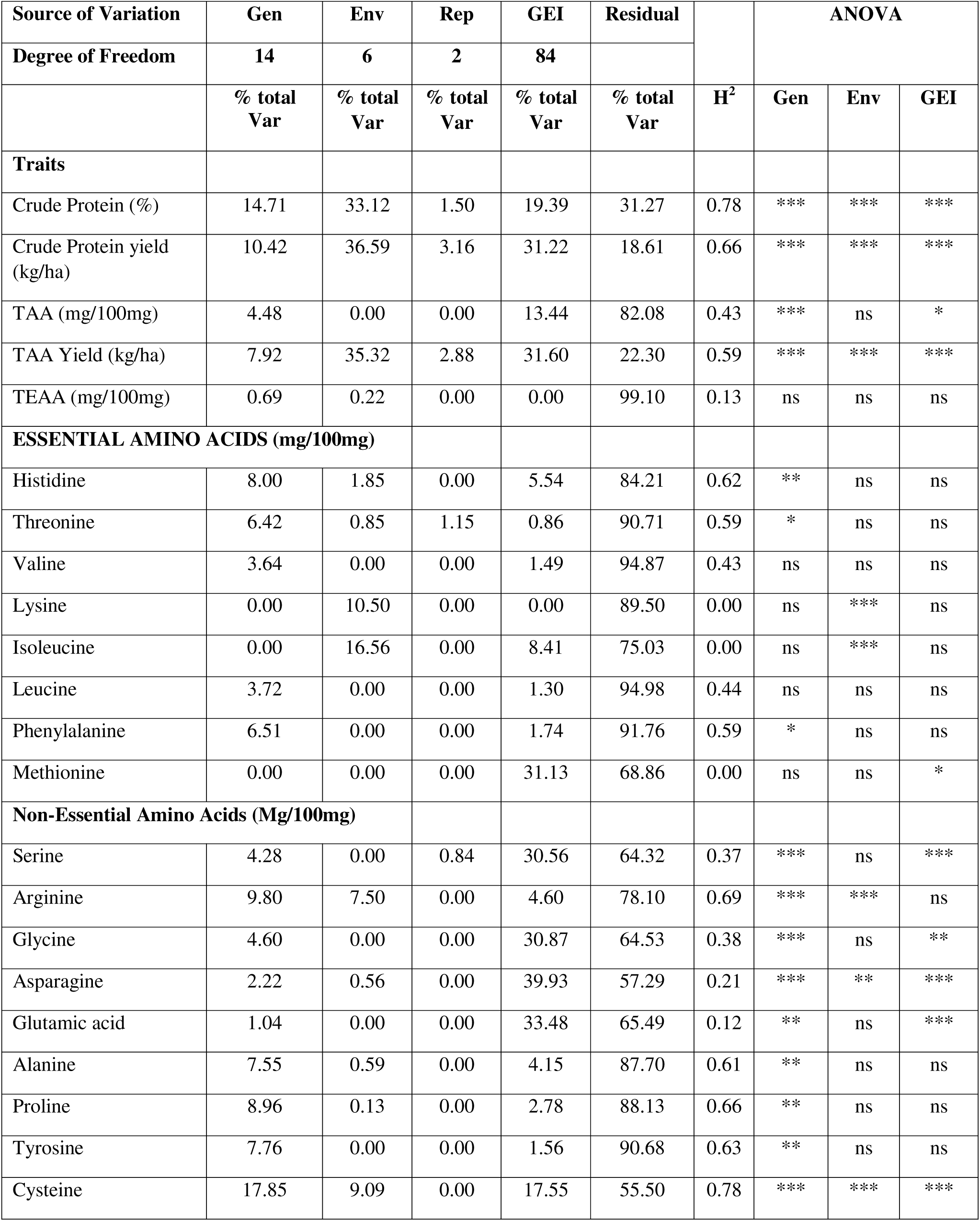
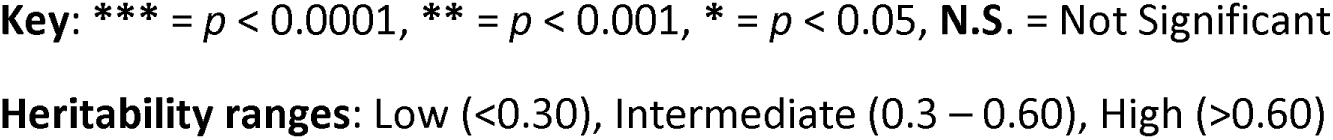
Variance partitioning among genotypic effect (Gen), environmental effect (Env), replication within the environment (rep), interaction genotype x environment (GEI), and residual (e). Broad-sense heritability (H^2^) is estimated from the regression between BLUP and phenotypic values and their respective ANOVA.

| Source of Variation | Gen | Env | Rep | GEI | Residual |  | ANOVA |  |  |
| --- | --- | --- | --- | --- | --- | --- | --- | --- | --- |
| Degree of Freedom | 14 | 6 | 2 | 84 |  |  |  |  |  |
| | % total Var | % total Var | % total Var | % total Var | % total Var | $H^2$ | Gen | Env | GEI |
| **Traits** |  |  |  |  |  |  |  |  |  |
| Crude Protein (%) | 14.71 | 33.12 | 1.50 | 19.39 | 31.27 | 0.78 | *** | *** | *** |
| Crude Protein yield (kg/ha) | 10.42 | 36.59 | 3.16 | 31.22 | 18.61 | 0.66 | *** | *** | *** |
| TAA (mg/100mg) | 4.48 | 0.00 | 0.00 | 13.44 | 82.08 | 0.43 | *** | ns | * |
| TAA Yield (kg/ha) | 7.92 | 35.32 | 2.88 | 31.60 | 22.30 | 0.59 | *** | *** | *** |
| TEAA (mg/100mg) | 0.69 | 0.22 | 0.00 | 0.00 | 99.10 | 0.13 | ns | ns | ns |
| **ESSENTIAL AMINO ACIDS (mg/100mg)** |  |  |  |  |  |  |  |  |  |
| Histidine | 8.00 | 1.85 | 0.00 | 5.54 | 84.21 | 0.62 | ** | ns | ns |
| Threonine | 6.42 | 0.85 | 1.15 | 0.86 | 90.71 | 0.59 | * | ns | ns |
| Valine | 3.64 | 0.00 | 0.00 | 1.49 | 94.87 | 0.43 | ns | ns | ns |
| Lysine | 0.00 | 10.50 | 0.00 | 0.00 | 89.50 | 0.00 | ns | *** | ns |
| Isoleucine | 0.00 | 16.56 | 0.00 | 8.41 | 75.03 | 0.00 | ns | *** | ns |
| Leucine | 3.72 | 0.00 | 0.00 | 1.30 | 94.98 | 0.44 | ns | ns | ns |
| Phenylalanine | 6.51 | 0.00 | 0.00 | 1.74 | 91.76 | 0.59 | * | ns | ns |
| Methionine | 0.00 | 0.00 | 0.00 | 31.13 | 68.86 | 0.00 | ns | ns | * |
| **Non-Essential Amino Acids (Mg/100mg)** |  |  |  |  |  |  |  |  |  |
| Serine | 4.28 | 0.00 | 0.84 | 30.56 | 64.32 | 0.37 | *** | ns | *** |
| Arginine | 9.80 | 7.50 | 0.00 | 4.60 | 78.10 | 0.69 | *** | *** | ns |
| Glycine | 4.60 | 0.00 | 0.00 | 30.87 | 64.53 | 0.38 | *** | ns | ** |
| Asparagine | 2.22 | 0.56 | 0.00 | 39.93 | 57.29 | 0.21 | *** | ** | *** |
| Glutamic acid | 1.04 | 0.00 | 0.00 | 33.48 | 65.49 | 0.12 | ** | ns | *** |
| Alanine | 7.55 | 0.59 | 0.00 | 4.15 | 87.70 | 0.61 | ** | ns | ns |
| Proline | 8.96 | 0.13 | 0.00 | 2.78 | 88.13 | 0.66 | ** | ns | ns |
| Tyrosine | 7.76 | 0.00 | 0.00 | 1.56 | 90.68 | 0.63 | ** | ns | ns |
| Cysteine | 17.85 | 9.09 | 0.00 | 17.55 | 55.50 | 0.78 | *** | *** | *** |

**Key:** \*\*\* = $p < 0.0001$ , \*\* = $p < 0.001$ , \* = $p < 0.05$ , **N.S.** = Not Significant
**Heritability ranges:** Low (<0.30), Intermediate (0.3 – 0.60), High (>0.60)

ANOVA showed a significant genotype effect for all the non-essential amino acids, environment effect was significant for Arg, Asp, and Cys and GEI effect were significant for Ser, Gly, Asp, and Glu (**Table 4**). Regarding the class of essential amino acids, genotype, environment, and GEI effects were not significant; considering specific EAA, significant effects were found for genotypes (His, Thr, and Phe), environment (Lys and Ile), and GEI (Met). However, the residual percentages of variation for individual amino acids showed high levels, indicating that a significant portion of the variance remained unexplained (**Table 4**). Genotype, environment, and GEI effects were significant for crude protein, crude protein yield, and TAAY, with most variance significantly attributable to the environment (**Table 4**). For TAA, a significant portion of the variance was due to GEI (13.4%) and genotype (4.5%), however, a high contribution was detected for residual (82.1%), indicating also in these cases that a significant portion of the variance is not explained, and the presence of other factors influences total amino acid level (**Table 4**).

Pearson correlation coefficients were computed among all traits (**Figure 2**). A significant and positive correlation was found between crude protein and TAA (r^2^ = 0.22, p = 0.0002). This correlation was also detected when performing the Simple Linear Regression between the CP and TAA by using single replicate data from all the field trials (**Supplementary Figure S5**). CP content shows a significant positive correlation with almost all the amino acids, except for Asp, Met, and Ile, while a negative and significant correlation was detected with Lys. Significant and positive correlations were detected among TAA and almost all the amino acids except for Lys and Ile. Comparing the different amino acid content across samples, we detected strong correlations among almost all of them (**Figure 2**). Ile and Lys did not correlate with any other amino acid. Met positively and significantly correlates with Asp, Gly, Pro, Ser, and Thr. Cys showed weak but significant and positive correlations with several amino acids, but not with Met, Lys, Tyr, and Ile.

**Figure 2.**
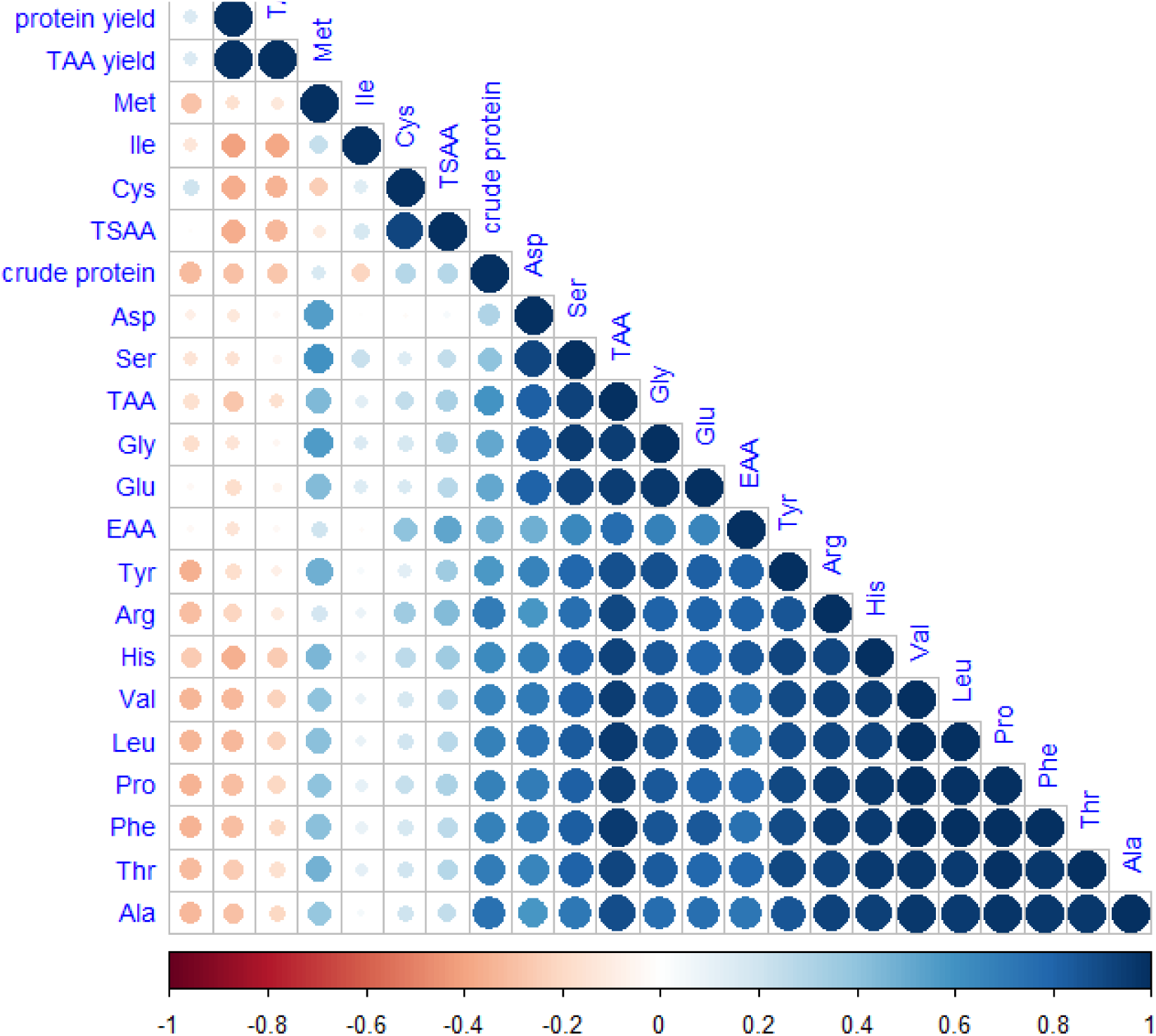
Pearson correlation between all traits.

### Phenotypic variability across genotypes

A Principal Component Analysis (PCA) representing the phenotypic space among the lentil genotypes for nutritional traits is provided in **Figure 3**. PC1 and PC2 accounted for 60.9% and 11.7% of the total phenotypic variance, respectively. PC1 allows separate genotypes based on the content of the majority of amino acids (except for Cys, Ile, Lys, and Met) and of crude protein, TAA, and TEAA (**Figure 3** and **Supplementary Table S3)**. PC2 was significantly and positively correlated with CPY and TAA yield, which are traits influenced by the nutritional composition and the overall yield, while it negatively correlates with Cys and TSAA content (**Supplementary Table 4**). The W627760 breeding line showed the highest CPY and TAA yield, while the PI431739 genotype showed the highest content of Cys and TSAA. The cultivars Flora, Anicia, and PI533693 were among the genotypes showing low content of AAs, CP, and TAA. We then compared genotypes, localities, and sowing seasons by performing an ANOVA, considering G, E, and GEI effects as fixed factors for CP, TAA, CPY, and TAAY (**Tables 5 and 6**; **Figure 4**). The analysis revealed that the effects of genotype (G), environment (E), and their interaction (G × E) were statistically significant for CP, CPY, and TAAY content. The influence of genotype (G) and the G × E had a significant impact on TAA composition, while a non-significant environmental effect was observed. CP content significantly varied among genotypes, with landraces showing the highest values, and in particular genotypes PI431710 and PI431739 (**Figure 4a**). Overall, a lower variability among genotypes was observed in TAA content; however, two landraces, PI431728 and PI431710, which are also the genotypes showing the highest CP content, exhibit the highest mean TAA content **(Figure 4b**). Regarding the TAA, landraces still exhibit the highest values, except for a breeding material (line PI_612875) that is among the best lines. Regarding CPY and TAAY, four genotypes (IG1959, PI431663, PI432033, PI612875) showed the highest values for both traits **(Figure 4b, d**).

**Figure 3.**
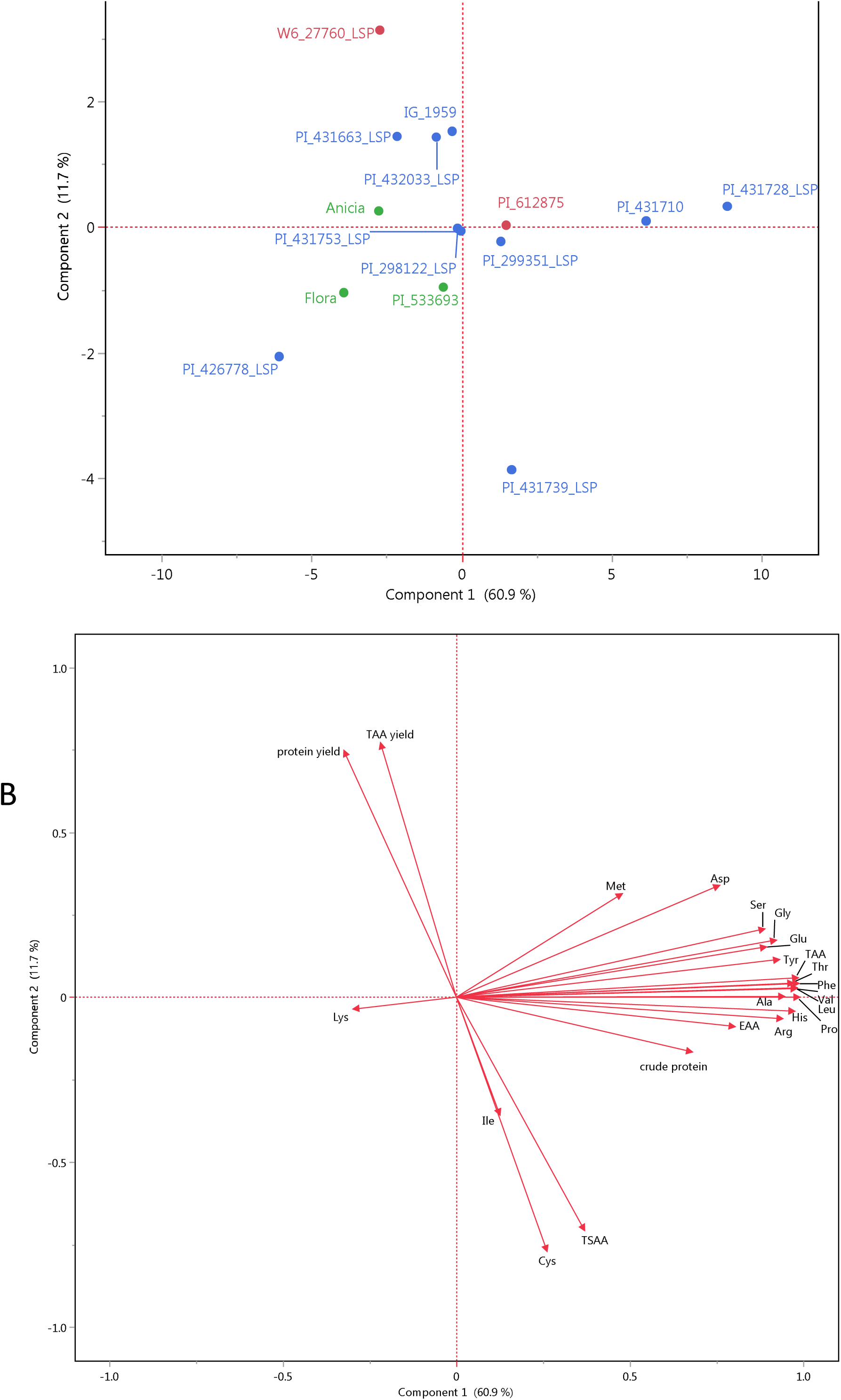
Principal component analysis based on the average values of all the traits for each genotype a) score plot highlighting the different biological status: landraces (blue), breeding material (red) and cultivar (green); **b)**the loading plot for all traits.PC1 and PC2 account for the 60.9% and the 11.7% of the total phenotypic variance.

**Figure 4.**
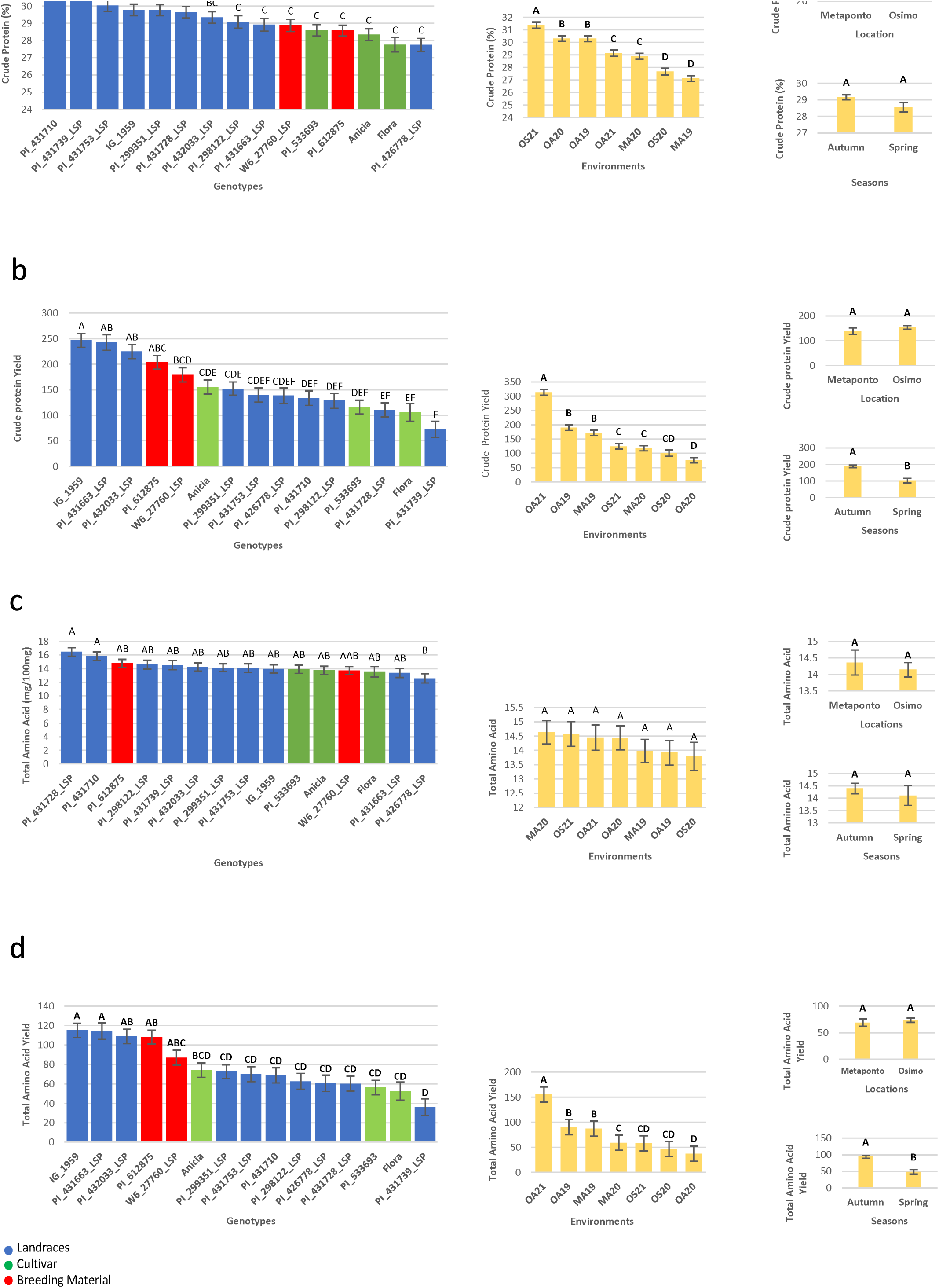
The influence of the genotype and the environments on the variation among CP, CPY, TAA and TAAY content for 15 lentil genotypes averaged across sites and years. (a)Least Square mean plot of seed CP(%)content for 15l entilgeno types averaged across 7 environments, alinear model with environments dissected into location and season effect for CP(%)content. (b)Least Square mean plot of seed CPY(kgha1)contents for 15l entil genotypes averaged across 7 environments, alinear model with environmnts dissected into location and season effect for CPY(kgha-1)content. ©Least Square mean plot of seed TAA(mg/100mg)content for 15l entil genotypes averaged across 7 environments, alinear model with environments dissected into location and season effect for TAA content. (d)Least Square mean plot of seed TAAY (kgha-1) content for 15l entil genotypes averaged across 7 environments, alinear model with environments dissected into location and season effect for TAAY(kgha-1) content.

**Table 5:** Analysis of variance for seed crude protein and crude protein yield for 15 lentil genotypes, considering all environments (years/location and sowing season).

| ANOVA |  | Crude protein |  |  | Crude Protein Yield |  |  |
| --- | --- | --- | --- | --- | --- | --- | --- |
| Source of Variation | df | F value | Pr (>F) | Proportion | F value | Pr (>F) | Proportion |
| ENV | 6 | 27.65 | *** | 33.12 | 20.72 | *** | 36.12 |
| GEN | 14 | 11.25 | *** | 14.71 | 17.97 | *** | 10.42 |
| GEN: ENV | 84 | 2.62 | *** | 19.39 | 5.98 | *** | 31.22 |

**Key:** \*\*\* = $p < 0.0001$ , \*\* = $p < 0.001$ , \* = $p < 0.05$ , **N.S.** = Not Significant

**Table 6:**
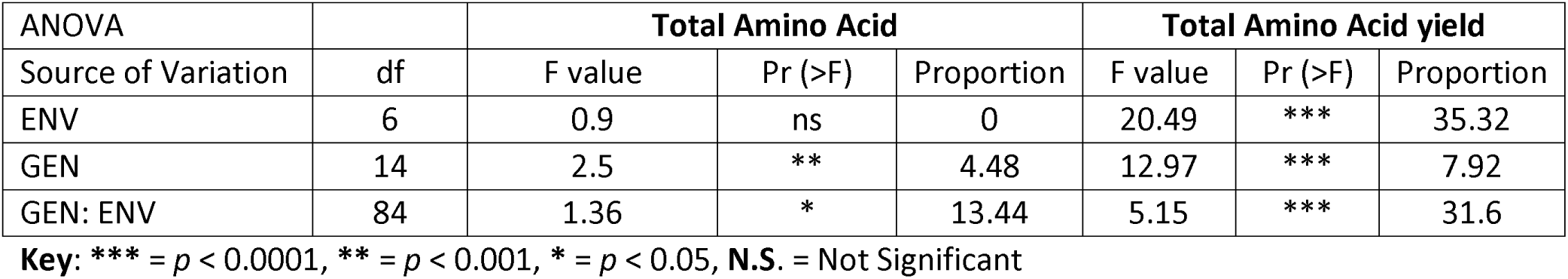
Analysis of variance for seed total amino acids and total amino acids yield for 15 lentil genotypes, considering all environments (years/location and sowing season).

No significant differences were detected among the seven environments for TAA content, nor localities and sowing seasons (**Figure 4c**). In contrast, significant variation was observed for CP content. In fact, the highest mean value was found in the OS21 field trial, followed by OA20 and OA19, while the lowest mean values were detected in OS20 and MA19 (**Figure 4a**). That is, while there was not a significant difference in terms of sowing season, we detected an overall higher production at the Osimo location. The OA21 field trial showed the highest nutritional yields, followed by OA19 and MA19; significantly higher mean values were observed in Osimo compared to Metaponto locality, as well as in autumn compared to spring sowing seasons (**Figure 4b, d**).

When we compared nutritional traits by grouping genotypes for biological status (landraces *versus* commercial and breeding varieties), only for crude proteins, we found that landraces are characterized by a significantly higher content (**Supplementary Figure S6**).

### Stability analysis of the genotypes for Crude Protein (CP) and Crude Protein Yield (CPY)

AMMI model-based stability analysis was conducted to partition genotype-by-environment interaction (GEI) effects for both CP content and CPY. For CP, the first and second interaction principal component axes (IPCAs) accounted for the majority of GEI variation, explaining 58.2% and 20.3% of the GEI sum of squares, respectively, capturing 78.5% of the total GEI (**Table 7**, **Figure 5a**). In the case of CPY, the first three IPCAs cumulatively explained 93% of the GEI variance (**Table 7**, **Figure 5b**).

**Figure 5.**
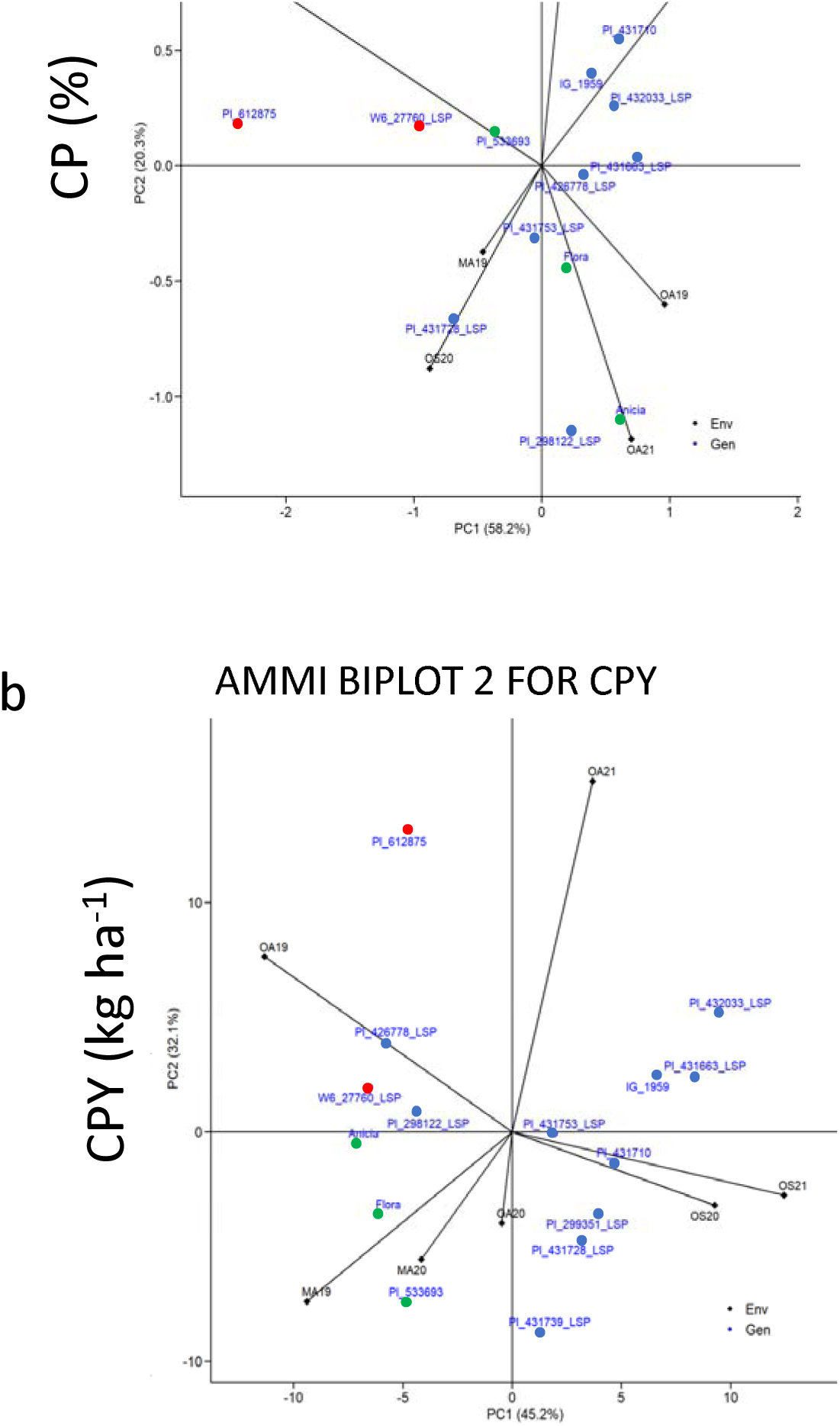
AMMI biplot representation of CP and CPY genotype × environment interaction. (a) AMMI2 biplot (PC1 vs PC2) for plotting CP data. (b) AMMI2 biplot (PC1 vs PC2) for plotting CPY data. Dots represent the 15 genotypes in common among all trials (Red, breeding material; blue, landraces; green, cultivars), while arrows and diamonds represent the seven environments, which are Osimo (OS) and Metaponto (MA) from 2019-2021.

**Table 7:** Additive main effects and multiplicative interaction (AMMI) results for seed crude protein and crude protein yield of 15 lentil genotypes, considering all environments (years, locations, and sowing seasons). The degree of freedom (Df), F value, and probability are reported in the table.

| Source | Df | Crude protein |  |  | Crude Protein Yield |  |  |
| --- | --- | --- | --- | --- | --- | --- | --- |
|  |  | F value | Pr<br>(>F) | Proportion (%) | F value | Pr<br>(>F) | Proportion (%) |
| **ENV** | 6 | 27.65 | *** | 33.12 | 20.72 | *** | 36.12 |
| **REP (ENV)** | 14 | 1.6 | ns | 1.5 | 4.36 | *** | 3.16 |
| **GEN** | 14 | 11.25 | *** | 14.71 | 17.97 | *** | 10.42 |
| **GEN: ENV** | 84 | 2.62 | *** | 19.39 | 5.98 | *** | 31.22 |
| **PC1** | 19 | 7.47 | *** | 58.2 | 13.12 | *** | 45.2 |
| **PC2** | 17 | 2.91 | *** | 20.3 | 10.42 | *** | 32.1 |
| **PC3** | 15 | 1.71 | ns | 10.5 | 5.76 | ** | 15.7 |
| **PC4** | 13 | 1.04 | ns | 5.6 | 1.52 | ns | 3.6 |
| **PC5** | 11 | 0.73 | ns | 3.3 | 1.43 | ns | 2.9 |
| **PC6** | 9 | 0.58 | ns | 2.1 | 0.33 | ns | 0.5 |
| **Residuals** | 165 |  |  |  |  |  |  |
| **Total** | 367 |  |  |  |  |  |  |

**Key:** \*\*\* = $p < 0.0001$ , \*\* = $p < 0.001$ , \* = $p < 0.05$ , **N.S.** = Not Significant

Two distinct environmental groups, Osimo Spring (2020 and 2021) and Metaponto Autumn (2019 and 2020), were particularly influential in driving GEI for both traits. Notably, lentil genotypes grown in Osimo Spring 2021 exhibited higher CP and CPY levels compared to those cultivated in Osimo Spring 2020. Conversely, the autumn environments were characterized by lower eigenvector values, indicating a reduced contribution to GEI relative to the spring sowing environments.

Most genotypes demonstrated limited interaction with the environment for both CP and CPY, including PI_431753_LSP, PI_431663_LSP, and PI_533693_LSP. In contrast, genotypes PI_612875, PI_431739_LSP, and Anicia exhibited significantly higher GEI effects, suggesting specific adaptation to particular environments (**Figure 5a**).

To integrate the AMMI analyses in the presence of a significant genotype-by-environment interaction (GEI) effect, genotype main effect (G) and genotype-environment (GGE) biplots for CP and CPY, commonly referred to as “which-won-where” plots, are presented in **Figure 6**. The combined contribution of the first two principal components (PC1 and PC2) to the variation in CP and CPY was 79.99% and 81.21%, respectively (**Figure 6**).

**Figure 6.**
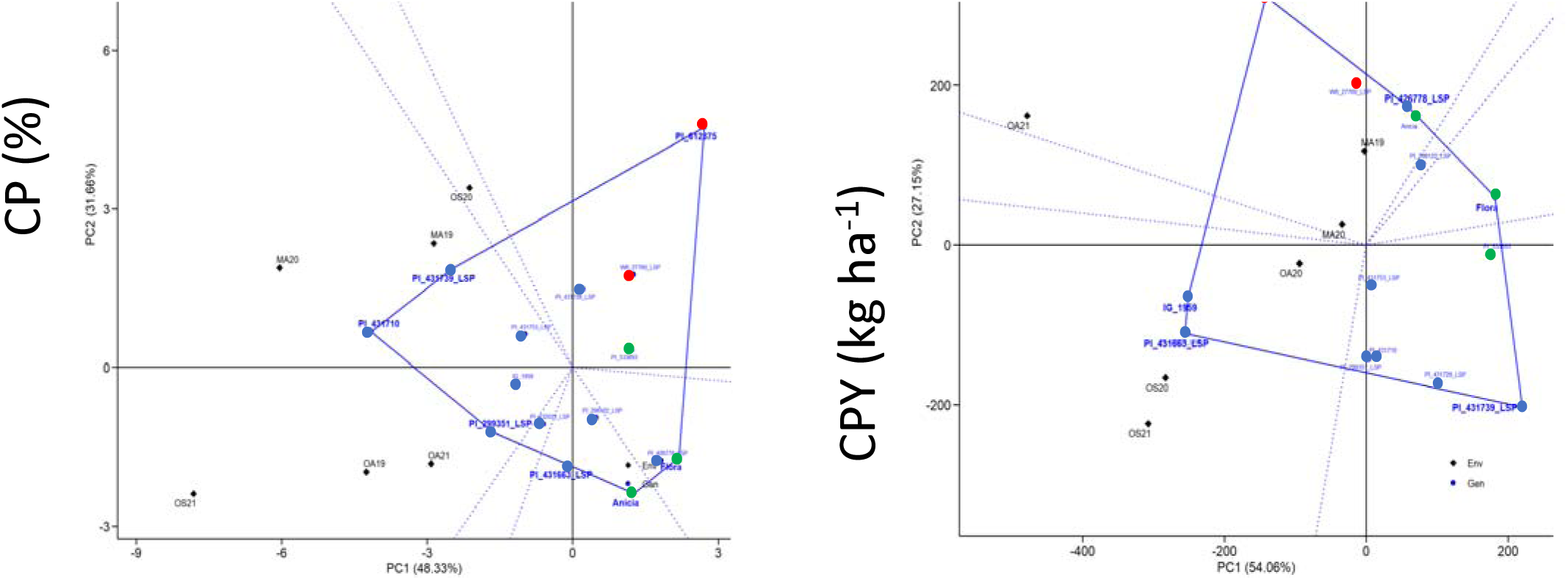
Polygon view of the GGE-biplot for the 15 lentil genotypes evaluated across seven environments in Osimo and Metaponto from 2019-2021 for the CP. **(a)** and CPY **(b)**. The “Which won where” GGE biplot. Blue dots represent the 15 genotypes in common among all trials and the green dots indicating different environments.

The biplot analysis for CP identified a mega-environment consisting of MA_19, MA_20, OA_19, OA_21, and OS_21, which predominantly represent autumn sowing seasons. Within this mega-environment, genotypes PI_299351_LSP, PI_431710, and PI_431739_LSP demonstrated superior performance (**Figure 6a**), while Flora, Itaca, PI_612875_LSP, and PI_431728_LSP showed the least favorable results. For CPY, the GGE biplot revealed two distinct mega-environments (**Figure 6b**). The first mega-environment, primarily encompassing the autumn growing seasons (MA_19, MA_20, OA_19, and OA_21), featured PI_612875 and PI_426778_LSP as the highest-performing genotypes. In contrast, the second mega-environment, which included both spring seasons (OS_20 and OS_21) as well as the OA_20 trial, highlighted IG_1959 and PI_431663_LSP as the top-ranking genotypes (**Figure 6b**).

Genotype stability was further assessed using the Y × WAASB biplot, with mean CP and CPY values plotted on the x-axis and WAASB values on the y-axis (**Figure 7**) as described by Olivotto (2019). Genotypes PI_431710, PI_431736_LSP, PI_431753_LSP, IG_1959, PI_431728_LSP, and PI_298122_LSP emerged as the most stable and productive for crude protein content, occupying the fourth quadrant that represents high performance and stability (**Figure 7a**). For CPY, IG_1959 and PI_431663_LSP were identified as the most stable and productive genotypes (**Figure 7b**).

**Figure 7.**
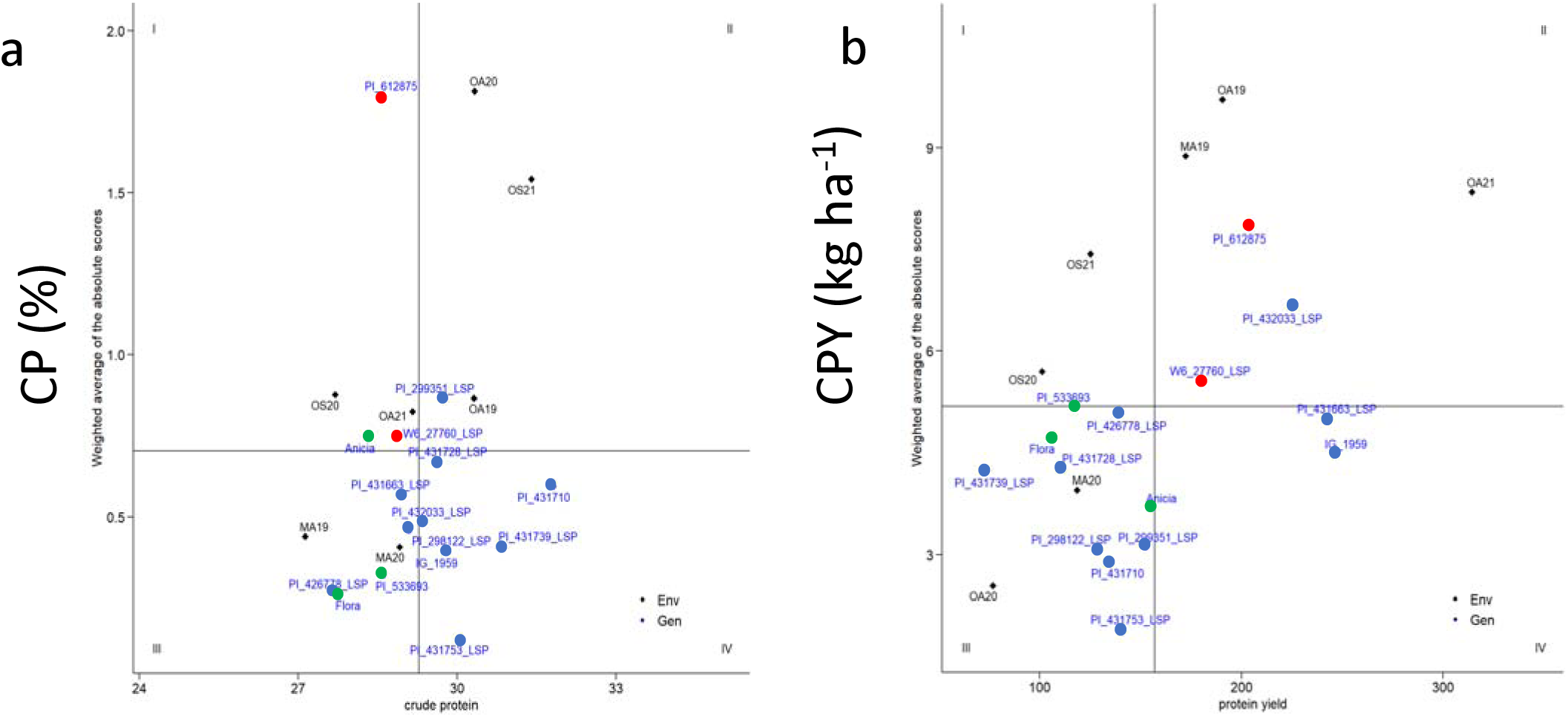
CP×WAASB (a) and CPY×WAASB (b) biplot based on joint interpretation of productivity (Y) and stability (WAASB) for 15 lentil genotypes evaluated under seven environments in Osimo and Metaponto from 2019-2021. Blue dots represent the 15 genotypes in common among all trials and the green dots indicating different environments. **Quadrant 1:** strong GEI component (high WAASBY values) and a below-average of response variable. **Quadrant 2:** strong GEI component (high WAASBY values) and an above-average of response variable. **Quadrant 3:** weak GEI component (stable) and a below-average of the response variable. **Quadrant 4:** weak GEI component(stable) and an above average of the response variable (recommended for breeding purposes.

### Stability analysis of the genotypes for Total amino acids (TAA) and Total amino acids yield (TAAY)

An AMMI model-based analysis was conducted to partition the genotype-by-environment interaction (GEI) for TAA. The results revealed that the first principal components significantly accounted for most of the observed variation, contributing cumulatively to 46.9% of the GEI (**Table 8**; **Figure 8a**). Similarly, for TAAY, the GEI was predominantly explained by the first three components, which together accounted for 94% of the variation (**Table 8**; **Figure 8b**). The AMMI biplot representation (**Figure 8**) highlighted two pairs of contrasting environments that primarily exemplified GEI for TAAY: Osimo Spring 2020 and 2021, as well as Metaponto Autumn 2019 and 2020 (**Figure 8b**).

**Figure 8.**
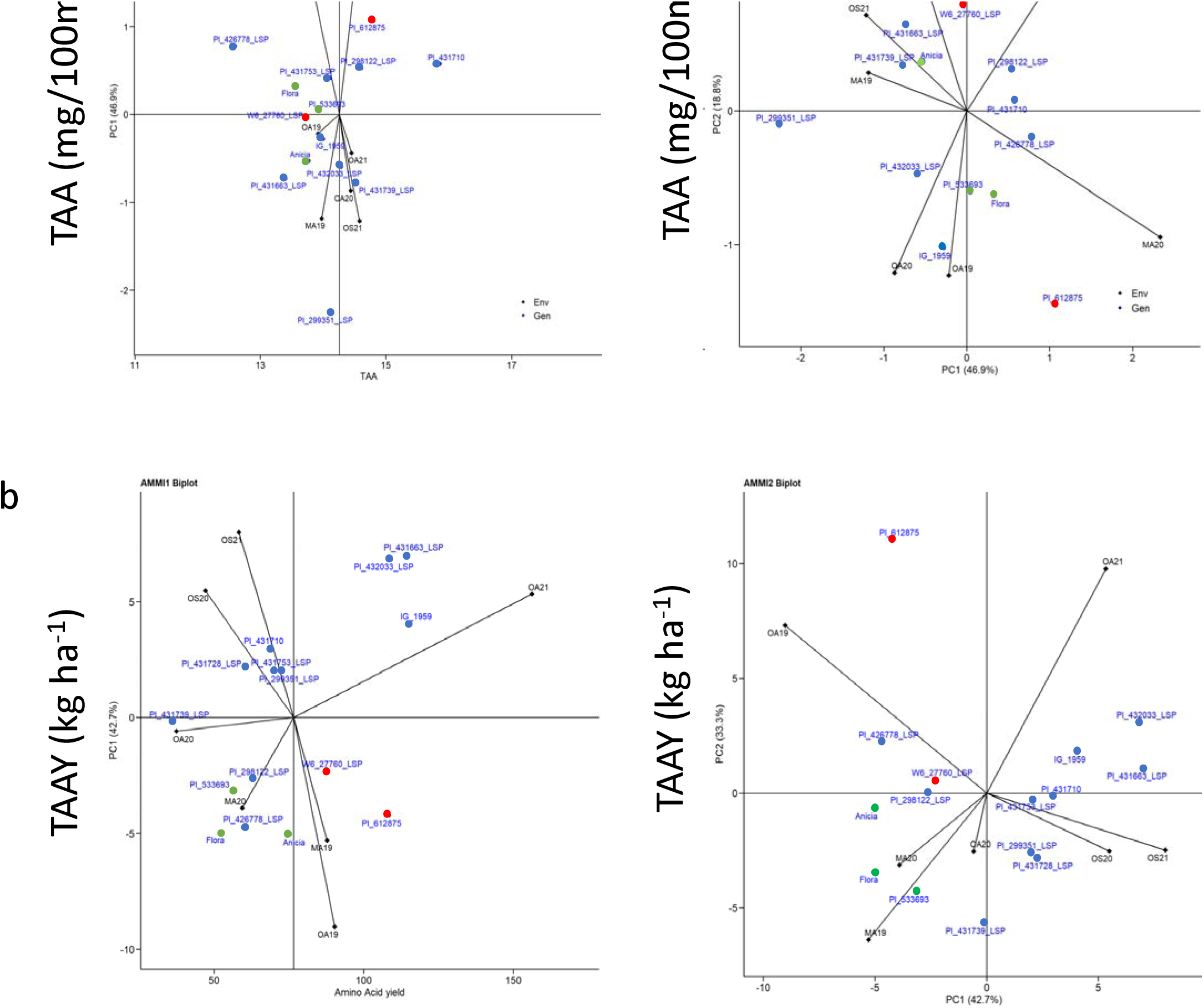
AMMI biplot representation of TAA and TAAY genotype × environment interaction. (a) AMMI1 biplot in the abscissa the genotype and environmental TAA and the ordinates the PC1 scores. (b) AMMI2 biplot (PC1 vs PC2) for plotting TAA data. Blue dots represent the 15 genotypes in common among all trials and the green dots the seven environments in Osimo and Metaponto from 2019-2021.

**Table 8.** Additive main effects and multiplicative interaction (AMMI) results for seed total amino acids and total amino acids yield for 15 lentil genotypes across environments and years.

| Source | Df | Total Amino Acids |  |  | Total Amino Acid Yield |  |  |
| --- | --- | --- | --- | --- | --- | --- | --- |
|  |  | F value | Pr (>F) | Proportion (%) | F value | Pr (>F) | Proportion (%) |
| ENV | 6 | 0.90 | ns | 0 | 20.49 | *** | 35.32 |
| REP(ENV) | 14 | 0.83 | ns | 0 | 3.49 | *** | 2.88 |
| GEN | 14 | 2.50 | ** | 4.48 | 12.97 | *** | 7.92 |
| GEN: ENV | 84 | 1.36 | * | 13.44 | 5.15 | *** | 31.6 |
| PC1 | 19 | 3.09 | ** | 46.9 | 10.56 | *** | 42.7 |
| PC2 | 17 | 1.39 | ns | 18.8 | 9.21 | *** | 33.3 |
| PC3 | 15 | 0.96 | ns | 11.5 | 5.65 | *** | 18 |
| PC4 | 13 | 0.92 | ns | 9.6 | 1.01 | ns | 2.8 |
| PC5 | 11 | 0.84 | ns | 7.4 | 1.02 | ns | 2.4 |
| PC6 | 9 | 0.81 | ns | 5.8 | 0.42 | ns | 0.8 |
| Residuals | 165 |  |  |  |  |  |  |
| Total | 367 |  |  |  |  |  |  |

**Key:** \*\*\* = $p < 0.0001$ , \*\* = $p < 0.001$ , \* = $p < 0.05$ , **N.S.** = Not Significant

Most of the tested genotypes, including PI_431710 and IG_1959, showed limited interaction with the environment, whereas PI_612875 and PI_426778_LSP were most affected by environmental variation (**Figure 8**).

The GGE biplot analysis for TAA and TAAY (**Figure 9**), illustrating the “which-won-where” pattern, indicated that the combined contribution of PC1 and PC2 to the variation in TAA and TAAY was 61.82% and 80.01%, respectively. For TAA, the biplot revealed two distinct mega-environments: the first included MA_20, OA_19, OA_21, and OS_20, with PI_431728_LSP and PI_426778_LSP emerging as the top-performing genotypes **(Figure 9a**). The second mega-environment, comprising MA_19, OS_21, and OA_20, identified PI_299351_LSP as the best-performing genotype. These mega-environments also exhibited a stochastic year effect (Figure 9a). The GGE analysis for TAAY (**Figure 9b**) produced results consistent with those observed in the CPY analysis (**Figure 9b**).

**Figure 9.**
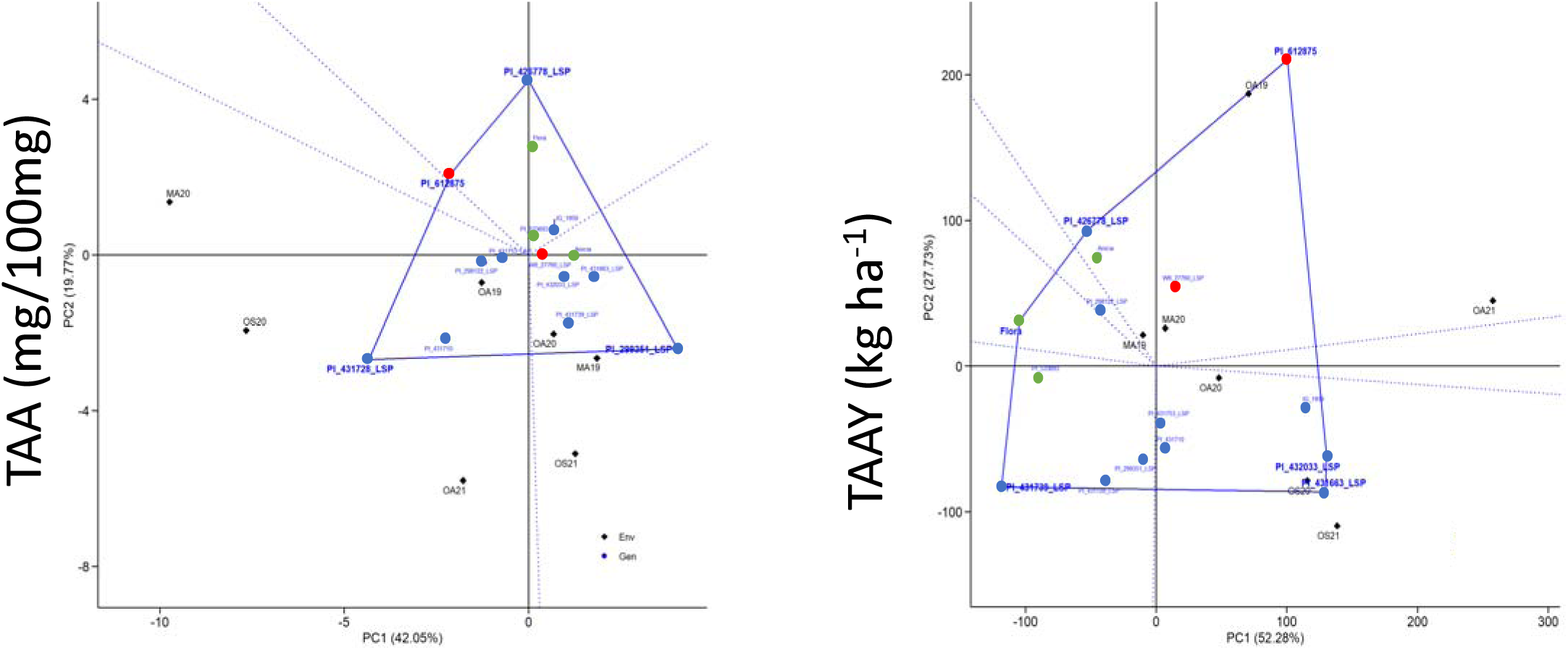
Polygon view of the GGE-biplot for the 15 lentil genotypes evaluated across seven environments in Osimo and Metaponto from 2019-2021 for the TAA (a) and TAAY (b). The “Which won where” GGE biplot. Blue dots represent the 15 genotypes in common among all trials and the green dots indicating different environments.

According to the WAABY biplot analysis, PI_431710, PI_298122_LSP, and PI_431739_LSP emerged as the most stable and high-performing genotypes for TAA content, as they were located in the fourth quadrant representing high performance and stability (**Figure 10a**). For TAAY, the genotypes W6_27760_LSP, PI_431663_LSP, and IG_1959 were identified as the most stable and highly performing (**Figure 10b**).

**Figure 10.**
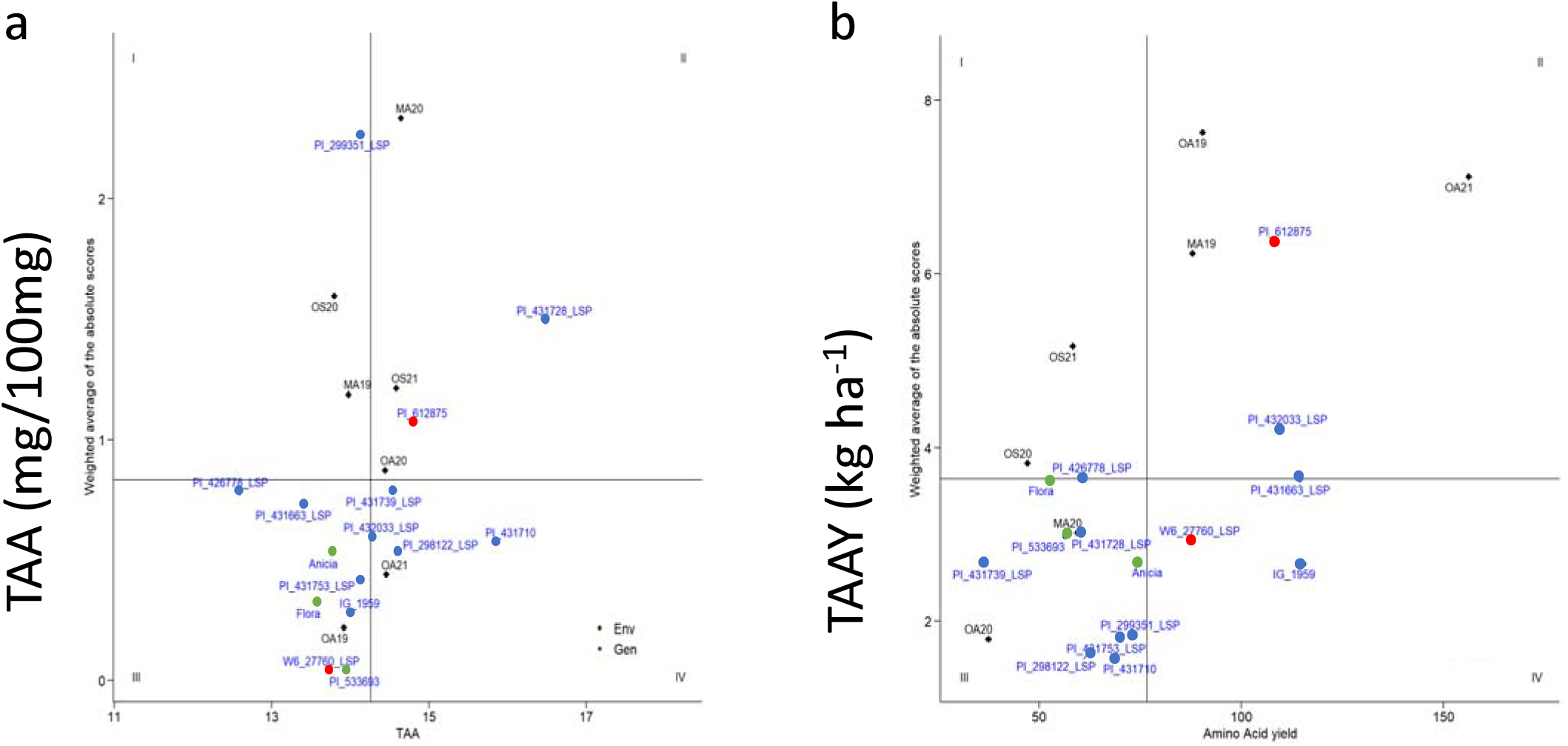
TAA×WAASB (a) and TAAY×WAASB (b) biplot based on joint interpretation of productivity (Y) and stability (WAASB) for 15 lentil genotypes evaluated under seven environments in Osimo and Metaponto from 2019-2021. Blue dots represent the 15 genotypes in common among all trials and the green dots indicating different environments. **Quadrant 1:** strong GEI component (high WAASBY values) and a below-average of response variable. **Quadrant 2:** strong GEI component (high WAASBY values) and an above average of response variable. **Quadrant 3**: weak GEI component (stable) and a below-average of the response variable. **Quadrant4**: weak GEI component(stable) and an above average of the response variable (recommended for breeding purposes).

## DISCUSSION

### Crude protein and total amino acid characterization of lentil genotypes

Lentil is a relevant crop for food in European and Mediterranean regions, which offer significant contributions to feeding the growing world population and agricultural sustainability (Khazaei et al., 2019; Montejano-Ramírez & Valencia-Cantero, 2024), being a valid alternative as a protein source (Bellucci et al., 2021). Lentils are recognized for their high protein content and balanced amino acid profile, making them an essential protein source in plant-based diets(Joshi et al., 2017). On one hand, it is crucial to identify novel genotypes that are well-adapted to different agro-environments, ensuring optimal yield and seed quality, to cope with the increasing demand for alternative proteins and the parallel climate change challenges. On the other hand, the limited breeding effort that can be recorded so far toward some of the most relevant legume species for food consumption is a major bottleneck for their cultivation and exploitation.

For these reasons, the agronomic characterization of genetic resources is crucial to assess new germplasm with important adaptive and yield traits, to be employed in breeding programs. In this study, we investigated the effect of genotype, environment, and their interactions (GEI) on key nutritional traits, such as crude protein (CP), total amino acids (TAA), and their respective yield components (i.e., crude protein yield [CPY] and total amino acids yield [TAAY]), that is the evaluation of seed quality combined with productivity. This work has been carried out considering 15 lentil genotypes of different biological status (i.e., landrace, cultivar, and breeding material), across seven field trials, including different sowing seasons (i.e., autumn and spring), years (2019-2020-2021), and locations (Osimo and Metaponto). Significant differences have been highlighted among genotypes, and environmental and GEI effects were detected. Thus, the study provides an important understanding of how lentil genetic diversity can be exploited for enhanced nutritional quality through targeted breeding programs (Baxevanos et al., 2024; Subedi et al., 2021; Yan et al., 2000). The amino acid composition of lentil seeds that has been described in this study aligns with previous research, highlighting relatively high levels of glutamic acid, arginine, and aspartic acid among non-essential amino acids (Ghumman et al., 2019; Khazaei et al., 2019; Paucean et al., 2018; Shekib et al., 1986). A deficiency in sulfur-containing amino acids (i.e., methionine and cysteine) was observed, in line with what is known from the literature (Carbonaro & Nucara, 2022; Khazaei et al., 2019; Salo-väänänen & Koivistoinen, 1996; Singh et al., 2023; Sree & Brar, 2020; Thavarajah et al., 2009). This emphasizes the need for dietary complementation with cereals, which are richer in these limited amino acids (Haug & Lantzsch, 1983; Messina, 2014; National Research Council, 1989; van Vliet et al., 2015). The significant genetic variation observed in non-essential amino acids (NEAAs) is of particular interest, as it supports the identification and selection of high-performing genotypes with stable expression across environments. This suggests strong potential for breeding improvements targeting these traits. In contrast, the environmental influence on essential amino acids such as lysine and isoleucine highlights the role of abiotic stressors like water availability and temperature in shaping nutritional profiles during critical developmental stages, including grain filling (S. Kumar et al., 2015; Margier et al., 2018).

Furthermore, our correlation analysis revealed a negative relationship between lysine and other nutritional traits, likely due to metabolic trade-offs in biosynthetic pathways (Angelovici et al., 2011; Yang et al., 2020; Zhu & Galili, 2004). Such trade-offs, where limited resources are allocated between growth and nutrient synthesis, are common in legumes and should be considered in future breeding strategies (Du et al., 2018). In addition, the study highlights the substantial influence of environmental factors on crude protein (CP) and crude protein yield (CPY) (M. S. Erskine, 1985; Gerrano et al., 2022; Subedi et al., 2021). While genotypic effects played a significant role in determining CP levels, CPY was more influenced by environmental variability in line with what has been previously suggested (Chen et al., 2022; Pugliese et al., 2024) and a significant GEI effect was also detected, particularly for CPY. Interestingly, landrace genotypes were always among the best performing genotypes when considering TAA, TAAY, CP and CPY; in particular, for CPY and TAAY, landraces IG_1959, PI_431663_LSP, and PI_432033_LSP, but also PI_431710 (for both CP and TAA), PI_431739_LSP (CP) and PI_431728 (TAA), are the most interesting lines, making them candidates for future breeding programs. The reduced protein content in modern cultivars suggests that past breeding efforts may have prioritized yield stability over nutritional quality, emphasizing the need for a balanced approach in future breeding programs (Sá et al., 2024). Likely, drought stress conditions recorded in Osimo Spring 2021 led to increased CP levels, a response often observed as plants redirect metabolic resources towards stress adaptation (Salehi, 2012; Zeroual et al., 2023). However, this came at the expense of seed yield and CPY, reinforcing the need for breeding strategies that balance protein content and environmental resilience (Bratković et al., 2024). Total amino acid (TAA) content and yield were primarily influenced by genotype and G × E interaction(Flores et al., 1998; Sellami et al., 2021; Shi et al., 2023). Environmental factors explained a significant portion of the variation in amino acid yield, underscoring the importance of environmental management in optimizing nutritional traits (Gerrano et al., 2022). Despite this, amino acid profiles remained relatively stable under moderate drought conditions, suggesting a degree of inherent resilience in lentil nutritional composition(Hang et al., 2022; Wu, 2013).

### Genotype by Environment Interaction (GEI) and Stability

In this work, we analyzed in more detail the GEI effect, in order to investigate relevant parameters such as genotype stability or adaptability to specific environments, to support plant germplasm selection, also for target environments.

Our analysis revealed that seasonal variations, particularly between autumn and spring plantings, were the primary drivers of GEI for both CP and CPY, confirming previous observations (Gauch Jr., 2006; Yan & Hunt, 1998). Among all the genotypes, landraces PI_431710_LSP, PI_431739_LSP, and PI_431753 were the most productive in terms of CP as stated above, and they can also be reported among the most stable; while IG1959 and PI_431663_LSP were the best performing and most stable with regards to the CPY. With regards to TAA, landrace PI_431728_LSP, that is the best performing genotype, was, however, less stable than other genotypes across environments, but still the genotype “who won” in the mega-environment, including three autumn sowing trials (MA20, OA21, and OA19) and OS20 field trial. On the other side, the second-best performing line for TAA, PI_43710 landrace, also proved to be stable across the considered environments. Finally, landrace IG1959 can also be considered as a promising material regarding the TAAY, based on the high performance and stability. Overall, our work highlighted a set of promising materials, considering stability, seed quality, and yield, but also including lines that outperformed in specific mega-environments, suggesting their potential exploitation as cultivated material or as being included in breeding programs for target environments. The identification of “mega-environments” in the GGE analysis further suggests that specific genotypes perform optimally under particular environmental conditions, providing valuable insights for region-specific breeding initiatives (Annicchiarico, 2002; Gauch Jr. et al., 2008). Of particular interest, landraces have been identified as among the most promising germplasm, making them a potential source of diversity and beneficial traits for seed quality, yield, and adaptability in modern lentil breeding and for direct cultivation (Ates, 2019; Babayeva et al., 2025; Lorenzetti et al., 2024). In our study, the energetic value of lentil genotypes over surface was not directly calculated in MegaJoule; literature shows that lentil cultivation efficiently produces substantial dietary energy, with reported outputs ranging from 10,000 to 15,000 MJ per hectare (approximately 2.4 to 3.6 million kcal/ha) (Mishra et al., 2017; Moraditochaee et al., 2014). In our work, however, we adopted the crude protein yield as a measure to evaluate the best-performing crop, highlighting the importance of considering what is usually classified as “quality” traits, such as nitrogen content and total protein in seeds, as crucial contributors to the overall yield. This is particularly relevant when considering the negative correlation that some crops and legumes, such as lentil, can display between the classical agronomic yield and the protein content, which does not allow a comprehensive evaluation of the overall quality of the energetic production. From the perspective of lentil breeding, we recommend integrating the overall protein yield with a more detailed analysis of the amino acid composition, for which we reported significant differences among genotypes.

### Conclusion

This study investigated and provided a significant understanding of the intricate dynamics between genotype, environment, and their interactions (G × E) in determining the nutritional quality of lentil (*L. culinaris* Medik.) seeds, particularly focusing on crude protein content and total amino acids (TAA). This research identified stable and high-performing genotypes and germplasm to be further exploited for breeding under target environments. The findings reveal that environmental factors play a significant role in determining protein yields, emphasizing the need for careful environmental management, alongside genetic selection. The use of advanced statistical models (AMMI, GGE, WAASB) provided a comprehensive understanding of these interactions, highlighting the complexity of breeding for stable nutritional traits in lentils. The identified stable genotypes offer promising avenues for breeding programs aimed at producing lentil varieties with enhanced protein content and amino acid profiles, which are crucial for meeting the dietary needs of growing populations. Future research should focus on integrating molecular and genomic tools to identify key genetic markers associated with protein content and amino acid profiles.

## Author contribution

V.D. conceived and managed the project; A.K.F. wrote the article; A.K.F., A.K., R.P., E.Bi, and V.D. contributed to the writing and drafting; A.K.F. and A. K performed data analysis; A.P., G.F., E.Be., L.N., L.R., C.S., E.Bi. contributed to the analyses; A.F.K., A.K., A.C., F.F., G.C., D.P., and B.F. conducted the nutritional profiling for crude protein and total amino acids; A.P., G.F., E.Be., T.G., L.N., L.R., C.S., D.P. contributed to the editing of the article; All authors read and approved the article.

## Supplementary figures

**Supplementary Figure S1:**
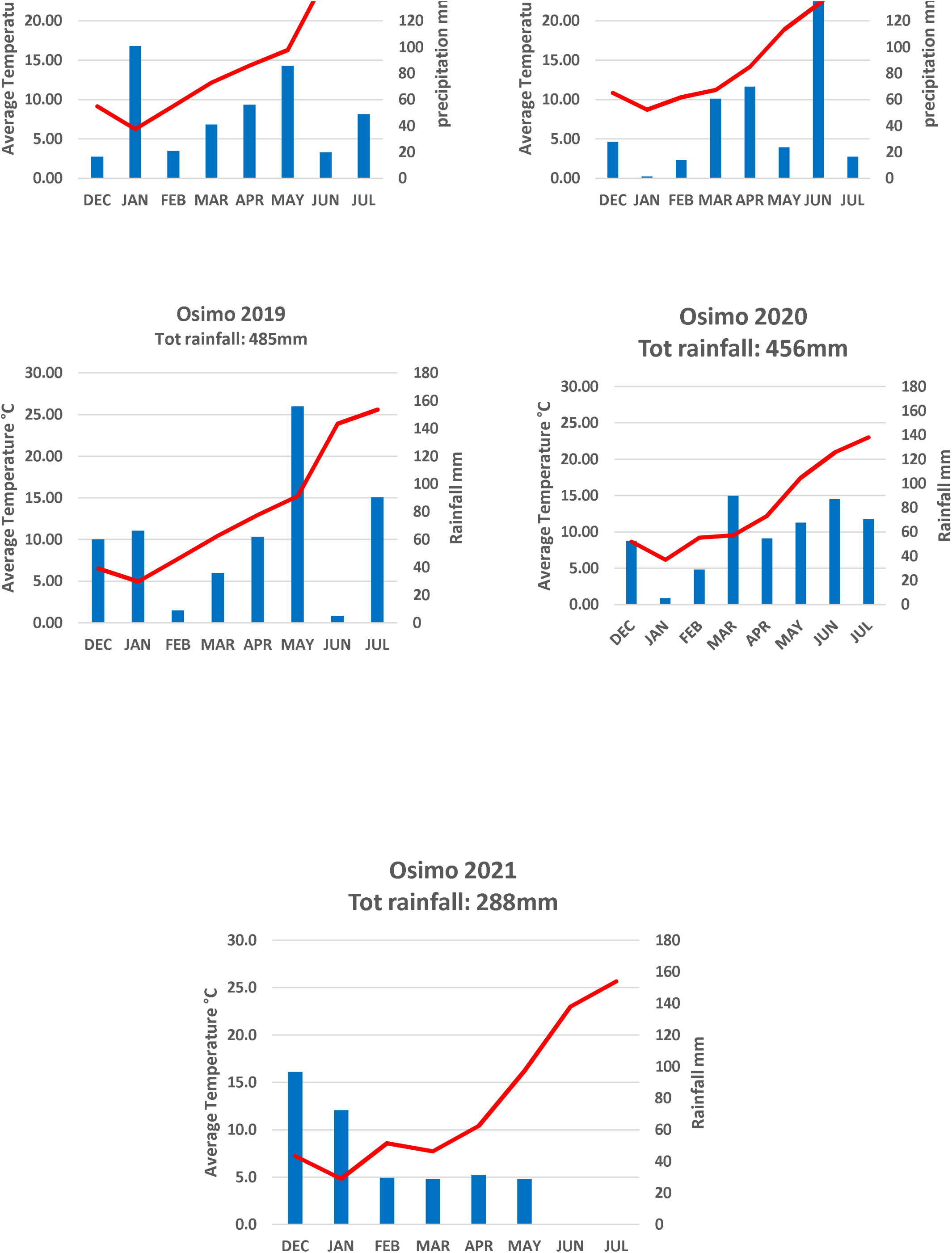
Thermo-pluviometric graph for the combinations of localities (Osimo, Metaponto) and years of trials (2019-2020-2021).

**Supplementary Figure S2.**
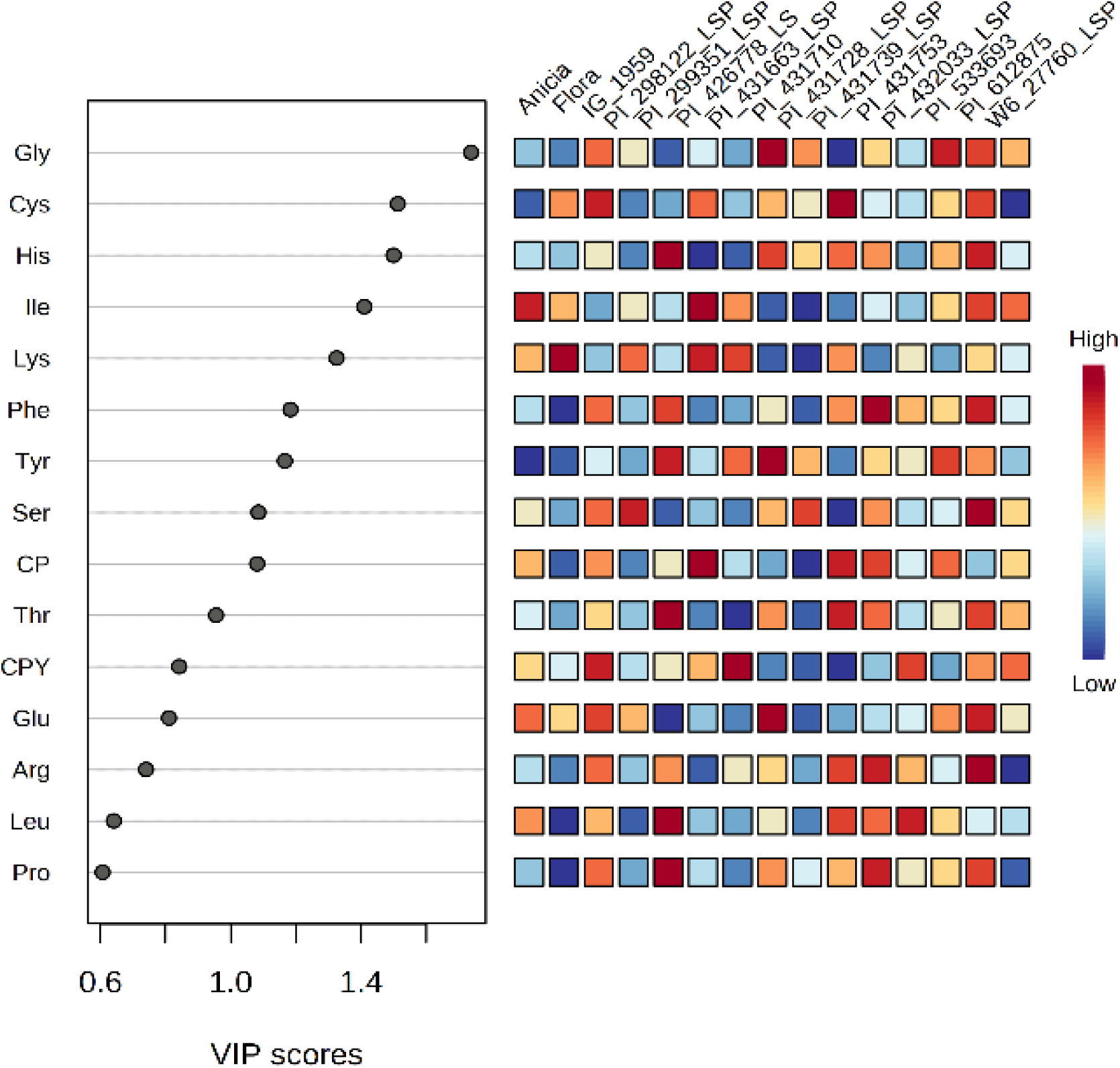
Variable Importance in Projection (VIP) plot.

**Supplementary Figure S3.**
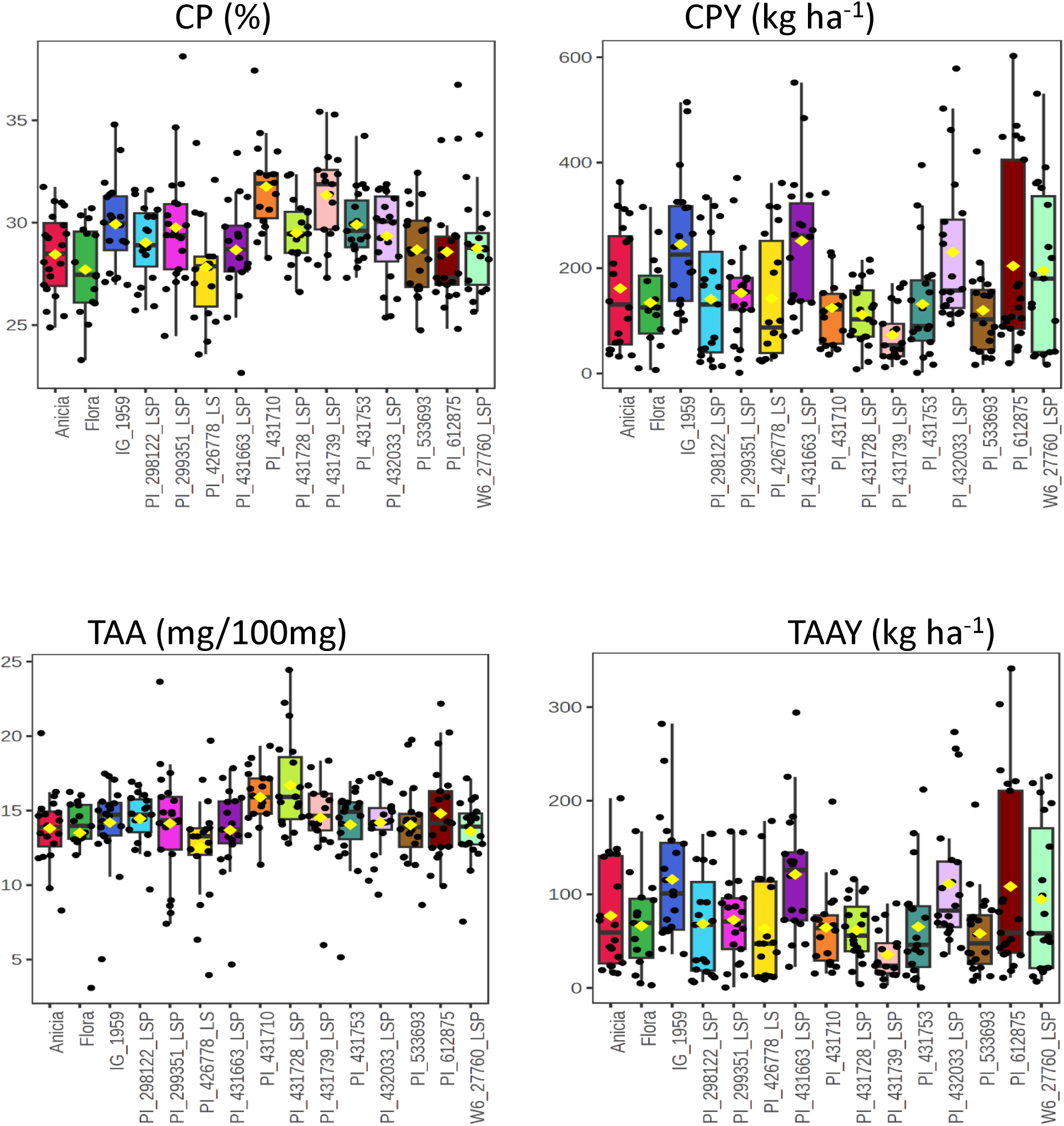
Representation of crude protein (CP), crude protein yield (CPY), total amino acid (TAA), and total amino acid yield (TAAY) profiles (nutritional quality) across 15 lentil genotypes. (a) Box plots of seed CP content (%), mean CPY (kg ha-1), TAA (mg/100mg), and TAAY (kg ha-1), content for 15 lentil genotypes averaged across sites and years.

**Supplementary Figure S4.**
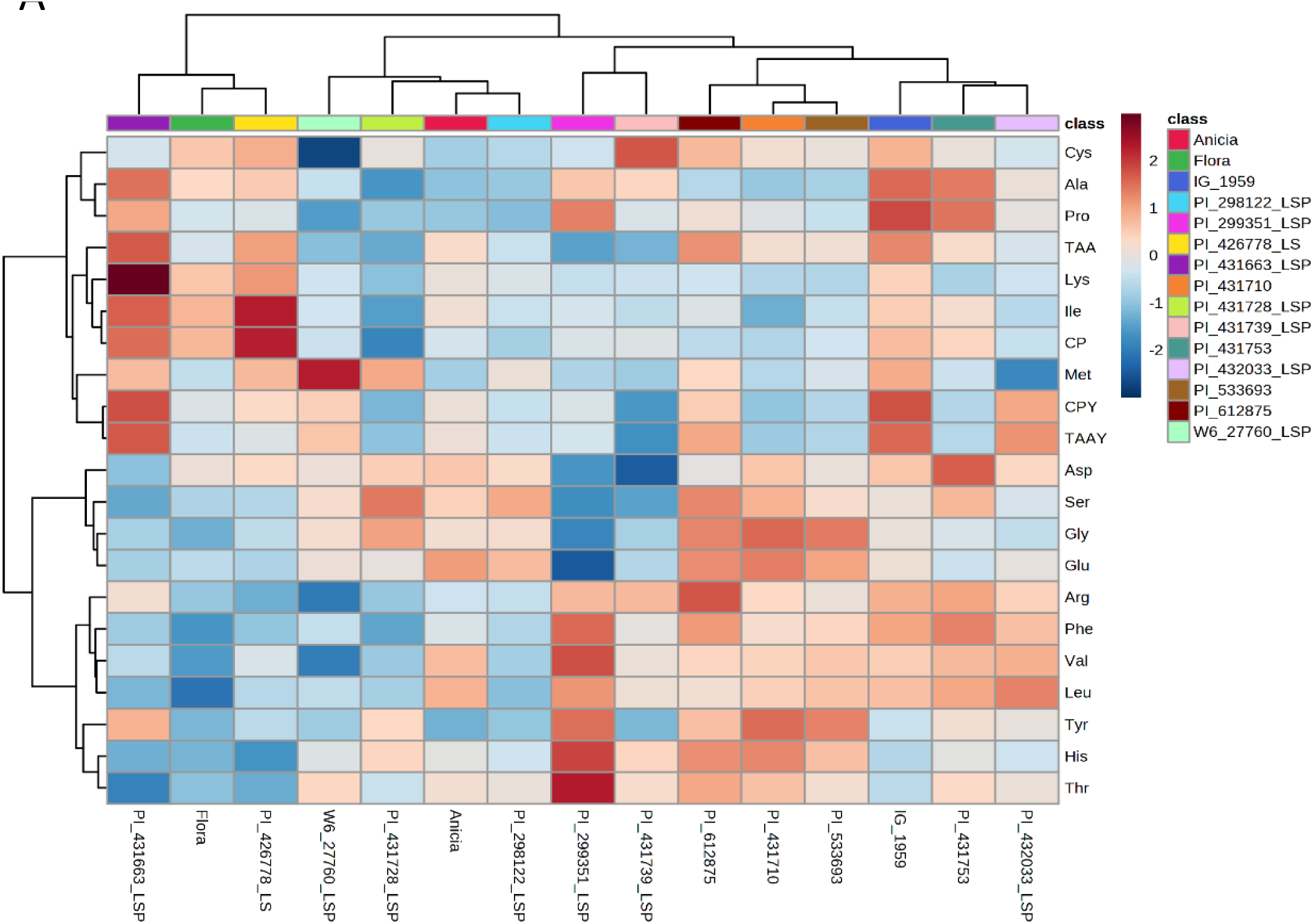
Representation of the crude protein (CP), crude protein yield (CPY), total amino acid (TAA) and total amino acid yield (TAAY) profiles (nutritional quality) across 15 lentil genotypes. (a) Heatmap reporting the average value of CP and of 17 AA profiling across all tested environments for 15 lentil genotypes. Regions of red or blue indicate lower or higher values compared to the average of each amino acids, respectively.

**Supplementary Figure S5:**
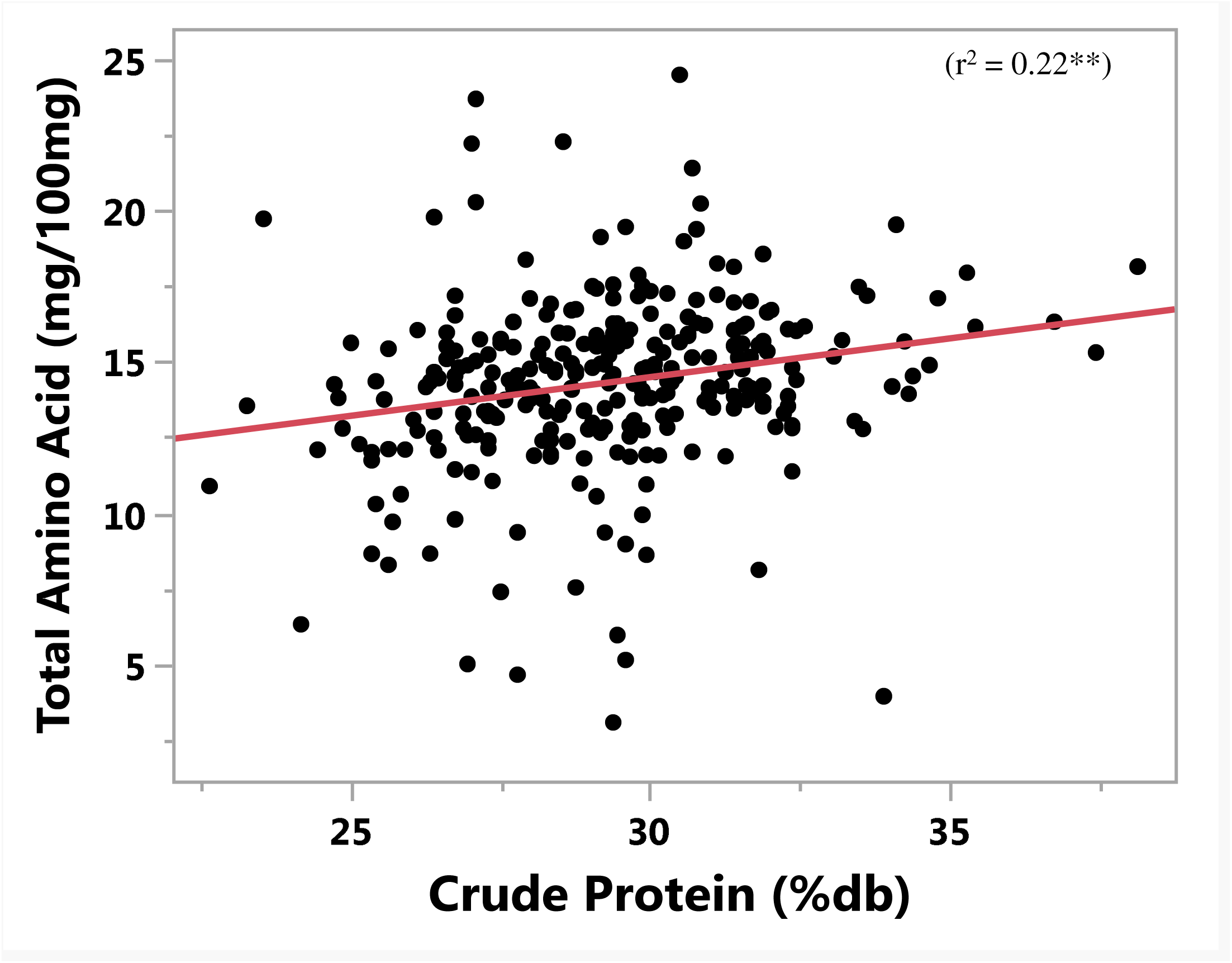
Linear Regression between the crude protein and TAA using single replicate data from all the field trials. **Key: \*\*** = p < 0.001.

**Supplementary Figure S6:**
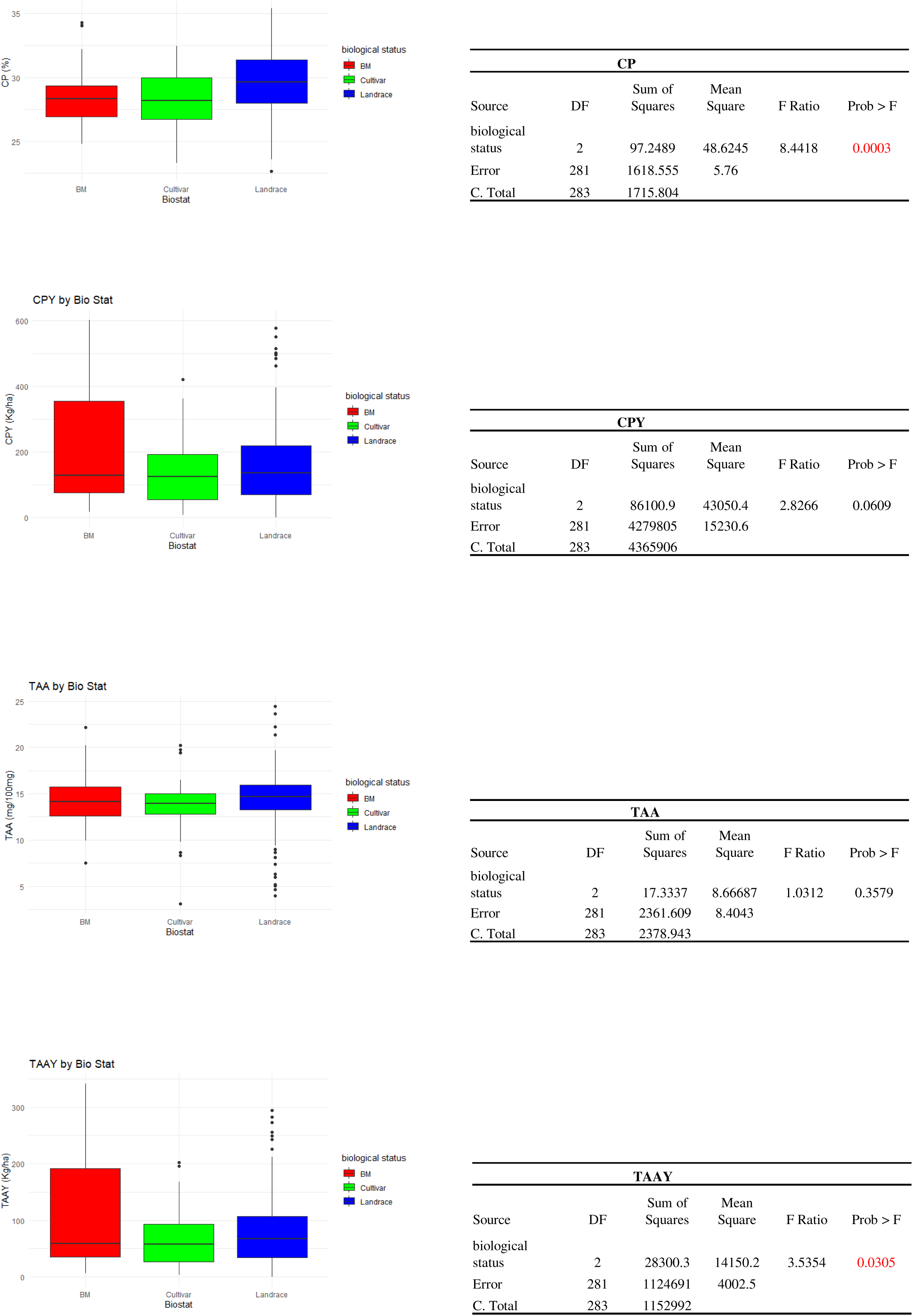
One-way ANOVA Traits and Biological statuses.

